# Single-cell mapping of tumor heterogeneity in pediatric rhabdomyosarcoma reveals developmental signatures with therapeutic relevance

**DOI:** 10.1101/2022.04.20.487706

**Authors:** Sara G Danielli, Ermelinda Porpiglia, Andrea J De Micheli, Natalia Navarro, Michael J Zellinger, Ingrid Bechtold, Samanta Kisele, Larissa Volken, Joana G Marques, Stephanie Kasper, Peter K Bode, Anton G Henssen, Dennis Gürgen, Josep Roma, Peter Bühlmann, Helen M Blau, Marco Wachtel, Beat W Schäfer

## Abstract

Rhabdomyosarcoma (RMS) is an aggressive human pediatric cancer. Despite robust expression of myogenic regulatory factors, RMS cells are blocked in a proliferative state and do not terminally differentiate. The extent to which the skeletal muscle lineage is represented in RMS tumors and the mechanisms leading to developmental arrest remain elusive. Here, we combined single-cell RNA sequencing (scRNAseq), mass cytometry (CyTOF) and high-content imaging to resolve RMS heterogeneity. ScRNAseq and CyTOF analysis of a total of 17 patient-derived primary cultures and three cell lines uncovered plastic myogenic subpopulations that delineate a branched trajectory. The less aggressive embryonal RMS (eRMS) harbor primarily muscle stem cell (MuSC)-like cells and exhibit sparse commitment to differentiation. The more aggressive alveolar RMS (aRMS) comprise primarily actively cycling committed progenitors with a paucity of differentiated cells. The oncogenic fusion protein PAX3:FOXO1 sustains aRMS cells in the cycling trajectory loop, which we show can re-wired towards differentiation upon its downregulation or by dual pharmacological RAF and MEK inhibition. Our findings provide insights into the developmental states and trajectories underlying RMS progression and identify the RAS pathway as a promising target of differentiation therapy for human aRMS.

**STATEMENT OF SIGNIFICANCE:** We present the first comprehensive single-cell transcriptomic and proteomic atlas of pediatric rhabdomyosarcoma (RMS), in which we identify impaired myogenic trajectories with prognostic value. We demonstrate that RAS pathway inhibitors disrupt the oncogenic trajectory and induce terminal differentiation, revealing novel therapeutic targets for the aggressive alveolar RMS subtype.

## INTRODUCTION

Intra-tumor heterogeneity (ITH) and phenotypic plasticity, new hallmarks of human cancer, pose challenges for the development of effective cancer therapies because they underlie drug resistance, tumor growth and metastasis formation^1, 2^. Understanding the mechanisms that govern these processes is crucial for designing more effective therapies that target the relevant cellular compartments. While childhood cancers mostly result from developmental defects in actively growing tissues^3^, emerging evidence suggests that both pediatric and adult tumors contain subpopulations of cells with different degrees of differentiation, that mirror the developmental stages of lineages found in the tissue of origin^4–7^. Therefore, characterizing the developmental hierarchies of individual tumors could be instrumental in identifying the cell of origin and in developing more effective treatments. A major limitation that has hindered progress to date has been the lack of tools to resolve tumor heterogeneity.

Rhabdomyosarcoma (RMS) is the most common pediatric soft tissue sarcoma and a highly aggressive cancer^8^. The two main RMS subtypes include embryonal RMS (eRMS) and alveolar RMS (aRMS). ARMS represents the more aggressive form of RMS and is mainly driven by a chromosomal translocation between the *PAX3* or *PAX7* and the *FOXO1* genes^8^. RMS tumors are believed to originate from differentiation defects during myogenesis, the process of skeletal muscle development^9^. Despite the expression of the transcription factors that drive myogenic commitment and differentiation, such as MyoD and Myogenin (MYOG), RMS cells are unable to complete myogenic differentiation, and therefore resemble undifferentiated progenitor cells. Previous reports proposed different states along the myogenic lineage as the candidate cells of origin, including mesenchymal progenitors^10^, satellite cells^11^ and mature differentiated myofibers^12^. More recent work implicated non-myogenic cells of origin, such as endothelial progenitor cells^13, 14^. However, the exact RMS cell of origin remains a matter of debate. Resolving the developmental heterogeneity and the mechanisms that lead to developmental arrest in RMS tumors is needed to define the cell origin definitively and inform therapeutic interventions.

In newly diagnosed RMS patients, one-third of localized and two-third of metastatic cases exhibit relapse or disease progression despite intense treatments^16^. In such cases, prognosis is dismal and 5-year overall survival rates drop below 10% for the aRMS subtype, highlighting the urgent need for therapeutic approaches^16^. While some eRMS tumors harbor druggable genetic alterations, the mutational landscape of aRMS tumors consists only of the oncogenic *PAX3/7:FOXO1* fusion itself^17, 18^. PAX3:FOXO1 is a potent and aberrant chimeric transcription factor that regulates proliferation, survival, and differentiation of aRMS cells, therefore representing an ideal, albeit challenging therapeutic target^19^. Recent findings suggest that PAX3:FOXO1 is dynamically expressed in a cell-cycle dependent manner and that its levels are heterogeneous within the aRMS cell population^20, 21^. Moreover, it remains unclear how PAX3:FOXO1 fluctuations influence subpopulation dynamics and whether aRMS cells retain their tumorigenicity after removal of PAX3:FOXO1^22^.

Differentiation therapy has emerged as a promising therapeutic strategy in diseases with dysregulated development^23, 24^. Despite the promises of a treatment that would push tumor cells to terminally mature and lose the malignant phenotype, clinical applications are still rare. Two recent studies demonstrated that myogenic differentiation can be triggered in *RAS*-mutant eRMS cells by interfering with RAS signaling^25, 26^, but differentiation of the more aggressive aRMS subtype has not been reported to date. A better understanding of the differentiation pathways in RMS is essential for designing effective differentiation therapies.

To tackle these challenges, we used a combination of single-cell RNA sequencing (scRNAseq), mass cytometry (CyTOF) and high-content imaging analysis to profile RMS cell lines and primary cultures derived from patient-derived xenografts (PDXs) that encompass the molecular and clinical diversity of human RMS. This approach allowed us to demonstrate that RMS tumors contain cell subpopulations that mirror the entire myogenic lineage. Using trajectory inference analysis, we built a differentiation model from single-cell transcriptomes to show that RMS muscle stem cell-like (MuSC-like) cells either differentiate or give rise to developmentally stalled actively cycling progenitors. We also show that primary cultures preserve broader potential for developing myogenic commitment and differentiation states compared to cell lines. We exploited this finding in a screen of pharmacological compounds to identify a combination of RAS pathway inhibitors, trametinib with dabrafenib or regorafenib, which triggers myogenic differentiation and cell cycle arrest in aRMS PDXs. Taken together, our findings provide a strong rationale for the clinical manipulation of the RAS pathway in aRMS patients and for inducing differentiation to counter oncogenicity.

## RESULTS

### Single-cell RNA sequencing of RMS PDX-derived primary cultures and cell lines identifies developmental heterogeneity in myogenic programs

To characterize the developmental traits of RMS tumors, we profiled 14 RMS PDX-derived primary cultures and 3 conventional aRMS cell lines by droplet-based scRNAseq (**Figure 1a**). To maximize inter-patient variability, we chose RMS models that originated from both aRMS and eRMS subtypes, primary and metastatic sites, diagnostic and recurrent patients that had or had not been pre-treated (**Extended Data Table 1**). After filtering for low quality cells, we retained 48,859 cells for downstream analysis. Cells clustered mainly by the patient of origin (**Figure 1b**), indicating substantial inter-tumor heterogeneity, a characteristic previously described for other tumor types^27^. To identify shared cell subpopulations and gene expression signatures, we integrated the RMS datasets using the “anchor” approach^28^. Unsupervised clustering resolved five distinct groups of cells that shared similar transcriptomic signatures (**Figure 1c; Extended Data Figure 1a; Supplementary Table 1**). We used gene set enrichment analysis^29^ in addition to published marker genes^30, 31^ (**Extended Data Table 2 and 3**), to group the cells into transcriptionally distinct cellular states. These cellular states included a novel early myogenic subpopulation (*MuSC-like*) which resembled muscle stem cells (MuSCs) and was enriched in vasculature development and cell adhesion genes; a proliferating subpopulation (*cycling progenitor*) that expressed high levels of cell division and cell cycle genes; two subpopulations of cells in G1 or S-phase that failed to express specific gene signatures, and a subpopulation of differentiated cells (*differentiated*), enriched in muscle system processes and myotube differentiation genes (**Extended Data Figure 1b and 1c**). We observed the same cellular states when integrating only aRMS or eRMS primary cultures or cell lines (**Extended Data Figure 1d; Extended Data Table 4**). The mesenchymal-associated markers CD44, ENG (CD105) and CXCL1 were mainly expressed by *MuSC-like* cells, the proliferation markers CDC20, CCNB1 and MKI67 by *cycling progenitors*, the committed muscle markers MYOG, ACTC1 and TNNT2 exhibited increased expression in cells undergoing differentiation and expression of myosin heavy (MyHCs) and light chain (e.g. MYH3, MYH8, MYL1) was exclusive of the *differentiated* subpopulation (**Figure 1d and 1e**). *MuSC-like* cells were the only aRMS subpopulation clearly characterized by absence of MYOG and by expression of CD44, PAX7 and AXL (**Extended Data Figure 1e**). To distinguish high-cycling from low-cycling cells, we scored each cell for validated gene signatures^32^. Notably, most *MuSC-like* and *differentiated* cells were low-cycling (**Figure 1f**), whereas cells in S-phase and *cycling progenitors* were high-cycling.

**Figure 1:**
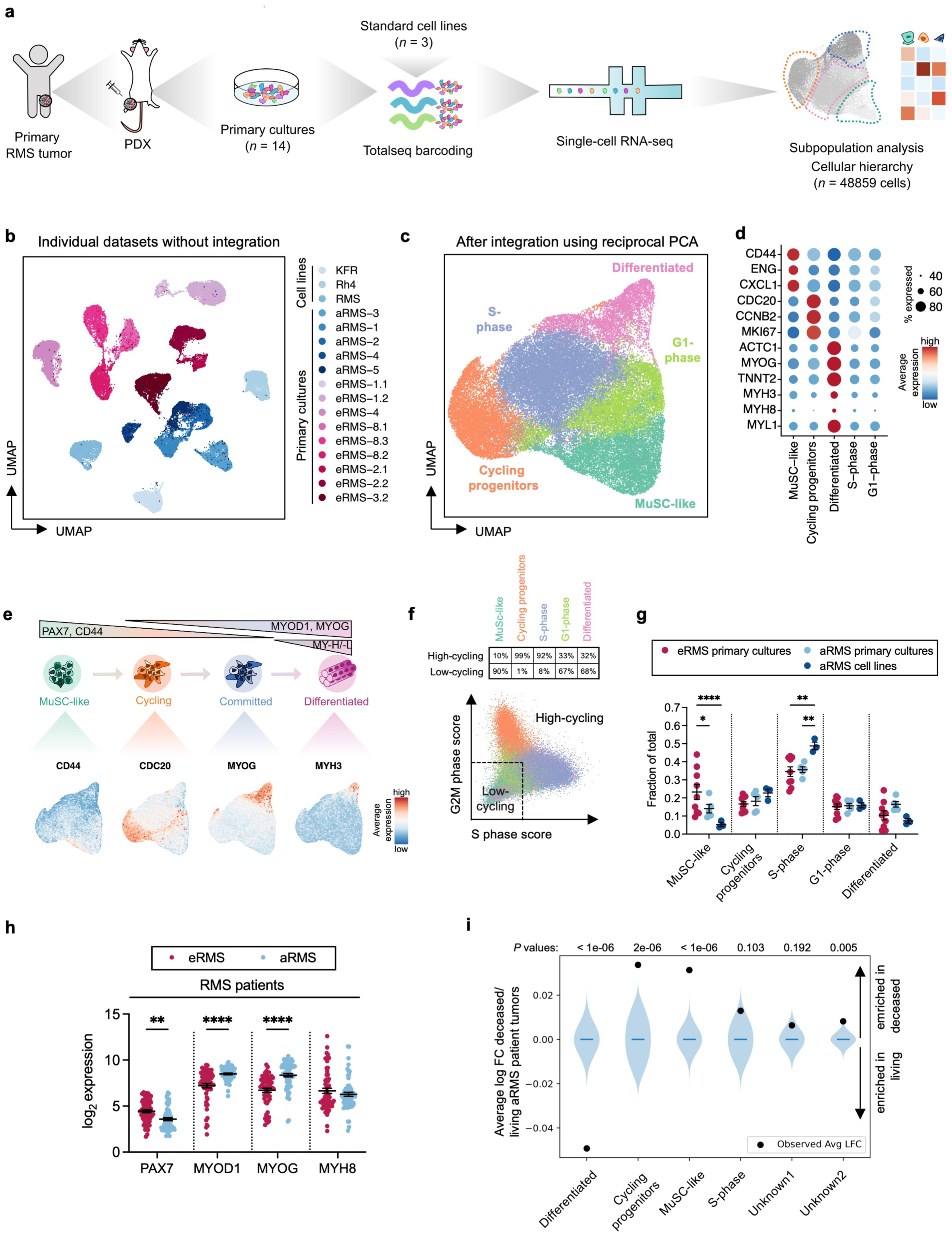
Single-cell RNA sequencing identifies heterogeneity in rhabdomyosarcoma primary cultures and cell lines recapitulating myogenic developmental programs. **a.** Experimental workflow. Patient’s tumors are first expanded in NSG mice as patient-derived xenografts (PDXs) and cultured for a low number of passages as primary cultures. Pools of n = 2 or n = 3 primary cultures were stained with Totalseq hashtags and loaded on 10X Chromium system for droplet-based scRNA-seq. **b.** UMAP plot of 48,859 cells from 14 RMS primary cultures and three cell lines after regressing the number of count RNA, the percentage of mitochondrial genes and the run batch effect. Cells are color-coded based on the corresponding sample of origin. **c.** UMAP plot of 48,859 cells after integration. Populations identified by Louvain clustering are shown. **d.** Dot plot showing expression of lineage-specific marker genes across the different Louvain clusters. **e.** Model of skeletal myogenesis with the populations identified in RMS. UMAP plots are colored based on the expression of markers delineating a myogenic lineage progression. **f.** Proportion of high-cycling (cells in G2M or S phase) and low-cycling RMS cells (cells in G1 phase) across the identified Louvain clusters. Cells are considered high-cycling if S-phase or G2M scores >0 and low-cycling if S-phase and G2M-phase scores <0. **g.** Relative proportion of Louvain clusters across RMS primary cultures and cell lines. Data are represented as mean ± SEM of *n* = 5 aRMS primary cultures, *n* = 3 aRMS cell lines and *n* = 9 eRMS primary cultures; ordinary two-way ANOVA with uncorrected Fisher’s LSD. **h.** Transcript levels of depicted markers in patient’s samples (*n* = 65 aRMS; *n* = 60 eRMS, R2 Davicioni dataset^33, 34^). Individual data points and mean ± SEM are shown; ordinary two-way ANOVA with uncorrected Fisher’s LSD. **i.** Association between cluster-associated signatures and aRMS patient (*n* = 65) survival. The aRMS patient dataset was downloaded from the R2: Genomics Analysis and Visualization Platform (http://r2.amc.nl; Davicioni -147 - MAS5.0 - u133a gene expression dataset). Violin plots show the log fold-change (FC) in the expression of the cluster-associated genes between deceased (status = dead) vs living patients (status = live) assuming there is no association (10^7^ simulation replicates). Black dots represent the calculated associations for each signature and the corresponding *P* values are listed. *, *P* < 0.05; **, *P* < 0.01; ****, *P* ≤ 0.0001.

Interestingly, cell lines mapped mostly to the *S-phase* cluster and had fewer cells in either *MuSC-like* or *differentiated* clusters compared to primary cultures; moreover, eRMS contained a higher proportion of cells mapping to *MuSC-like* states than aRMS (**Figure 1g**). We confirmed these differences in a large RMS patient mRNA expression dataset available on the R2 platform^33, 34^. Compared to eRMS, aRMS patients showed significantly lower PAX7, known to mark muscle stem cells, higher MYOD1 and MYOG, known to mark activated and committed cells, and no difference in MYH8 mRNA expression, known to mark differentiated myogenic cells (**Figure 1h**). Finally, we evaluated the clinical impact of these transcriptional signatures by testing whether they were associated with different clinical outcomes in aRMS patients^34^. Strikingly, the *MuSC-like* and *cycling progenitor* signatures were highly represented in deceased aRMS patients, whereas the *differentiated* signature was strongly enriched in living aRMS patients (**Figure 1i**). Both these associations revealed to be strongly statistically significant. All together, these findings support the notion that RMS tumors contain myogenic cells stalled in an immature transcriptional state, but they reveal a cellular continuum that spans the entire myogenic lineage from *MuSC-like* to a minority of *differentiated* cells. They also identify an intermediate *cycling progenitor* subpopulation that correlates with worst outcomes in aRMS patients and a subpopulation resembling differentiated muscle cells, which correlates with better outcomes.

### Single-cell mass cytometry reveals novel RMS subpopulations that delineate a distinct myogenic progression

To confirm the presence of myogenic subpopulations at the protein level, we profiled RMS primary cultures by mass cytometry (CyTOF). We pulsed cells *in vitro* with iododeoxyuridine (IdU), to determine their cell cycle status, and stained them with an isotope-conjugated antibody panel, which included known surface markers used to isolate MuSCs, novel surface markers identified by scRNAseq, and myogenic transcription factors known to define distinct stages of myogenesis^35^. We analyzed the pooled aRMS and eRMS CyTOF datasets with the clustering algorithm X-shift^36^, using novel surface markers (CD44, Axl), myogenic transcription factors (Pax-7, myogenin) and a cell cycle marker (IdU) (**Figure 2a**). We visualized the spatial relationship between the resulting clusters by single-cell force-directed layout, to build a cell-cycle driven map of RMS in which the physical distance between clusters indicates their similarity in marker expression in high dimensional phenotypic space. The map was then colored based on individual markers, including the myogenic transcription factors Pax-7, MyoD and myogenin which delineate a myogenic progression from muscle stem cells to activated and committed/differentiated myogenic cells, respectively. Data analysis revealed that while cells from aRMS and eRMS shared some commonalities, they also occupied unique regions that were specific to each tumor (**Figure 2b**). We defined cells that were actively incorporating IdU, a marker of S-phase, as high-cycling cells^37^, whereas cells that were not incorporating IdU were defined as low-cycling. Pax-7^+^ cells were present in both subsets, which led to identification of *quiescent MuSC-like* cells and *proliferating MuSC-like* cells in both aRMS and eRMS. However, Pax-7 expression was higher in eRMS cells and IdU incorporation was higher in aRMS cells. These findings are in line with the scRNAseq analysis described above. A unique signature of eRMS revealed by our CyTOF analysis includes a cell subset that expresses high levels of MyoD. On the other hand, unique signatures of aRMS include two prominent myogenin^hi^ subsets, one of which incorporated high levels of IdU, and was therefore cycling (*cycling progenitors*). This subset is of particular interest because it represents a population of cells that has never been observed in myogenesis in adult skeletal muscles: it proliferates and at the same time expresses the differentiation marker myogenin, which under normal circumstances causes differentiation and cell cycle exit. Our findings suggest that the newly identified myogenin^hi^/IdU^hi^ aRMS subset is defective in inducing the myogenic differentiation program. Hence, this novel subset could serve as a cellular target for RMS differentiation therapy. In summary, we identified four developmental stages in RMS primary cultures, which include *quiescent MuSC-like* cells (IdU^-^/Pax-7^+^/myogenin^-^), *cycling MuSC-like cells* (IdU^+^/Pax-7^+^/myogenin^-^), *cycling progenitors* defective in differentiation (IdU^+^/myogenin^+^) and *non-cycling committed/differentiated progenitors* (IdU^-^ /myogenin^+^). Importantly, we found that co-expression of the cell surface proteins CD44, AXL and CD105 marked the aRMS *MuSC-like* cell subpopulation. This finding provides a strategy to prospectively isolate a cell subpopulation characteristic of aRMS for in depth functional studies.

**Figure 2:**
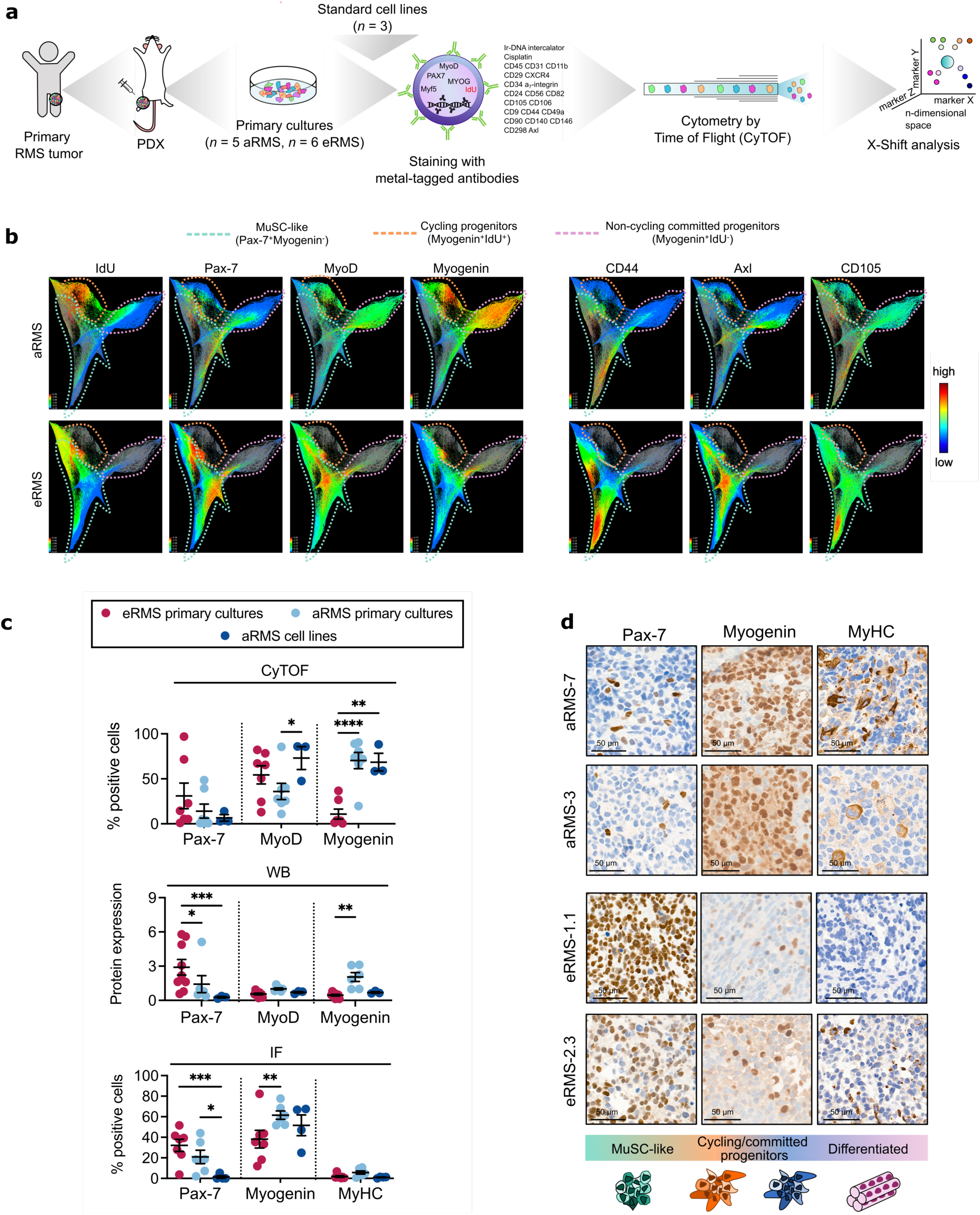
Single-cell protein profiling defines RMS subpopulations with high-resolution. **a.** CyTOF mass cytometry workflow. Primary cultures and cell lines were stained with isotope-conjugated antibodies, including known and novel cell surface markers, and myogenic transcription factors, and run through the CyTOF instrument. Data were analyzed using the clustering algorithm X-shift. **b.** Human Live/CD298^+^ cells gated from *n* = 11 primary cultures (pooled aRMS and pooled eRMS) were clustered with the X-shift algorithm. The resulting clusters were visualized using single-cell force-directed layout, to generate a cell-cycle driven map of RMS (Samusik et al., *Nature Methods*, 2016). The cells on the map are colored by IdU incorporation, to mark proliferating cells, or by marker expression (aRMS top panel, eRMS lower panel). The cell surface markers CD44, Axl and CD105 mainly mark aRMS cells in the *MuSC-like* cluster. **c.** Percentage of positive cells and expression levels of markers delineating a myogenic progression (Pax-7, *MuSC-like* cells; MyoD, activated cells; myogenin, committed/differentiated cells; MyHC, *differentiated* cells) across individual RMS primary cultures and cell lines measured by CyTOF (top panel; *n* = 5 aRMS primary cultures, *n* = 6 eRMS primary cultures, *n* = 3 aRMS cell lines), western blot, WB (intermediate panel; *n* = 6 aRMS primary cultures, *n* = 9 eRMS primary cultures, *n* = 3 aRMS cell lines) and immunofluorescence, IF (lower panel; *n* = 6 aRMS primary cultures, *n* = 7 eRMS primary cultures, *n* = 4 aRMS cell lines). Data are represented as mean ± SEM of the indicated number of samples; ordinary two-way ANOVA with uncorrected Fisher’s LSD. *, *P* < 0.05; **, *P* < 0.01; ***, *P* < 0.001; ****, *P* ≤ 0.0001 **d.** Expression of Pax-7, myogenin and MyHC markers as determined by immunohistochemistry in PDX tumors. *, *P* < 0.05; **, *P* < 0.01; ***, *P* < 0.001; ****, *P* ≤ 0.0001

To confirm the differential cellular distribution of aRMS and eRMS subtypes across myogenic stages, we measured the proportion of cells arrested at the progenitor (Pax-7^+^), activated (MyoD^+^), committed (myogenin^+^) and differentiated stages (MyHC^+^) in several primary cultures and cell lines by CyTOF, immunofluorescence (**Extended Data Figure 2a**) and/or western blot analysis (**Extended Data Figure 2b**). ARMS primary cultures comprised a larger fraction of myogenin^+^ cells and expressed significantly lower Pax-7 but higher myogenin levels compared to eRMS; cell lines comprised a significantly smaller fraction of Pax-7^+^ cells compared to primary cultures (**Figure 2c**), confirming the more homogeneous composition of the cell lines, which was also found in the scRNAseq clustering distribution. To demonstrate the presence of these subpopulations at the tissue level we performed immunohistochemistry (IHC) analysis of solid PDX tumor sections. We uncovered heterogeneity in the expression levels of Pax-7, myogenin and MyHC. Importantly, we found that aRMS PDXs comprised a smaller number of Pax-7^+^ cells and larger number of myogenin^+^ and MyHC^+^ cells (**Figure 2d**). Altogether, our single-cell protein analysis corroborates the transcriptomic analysis.

### Single-cell trajectory inference uncovers a branched lineage organization of RMS primary cultures

We investigated the relationship between different RMS subpopulations using a trajectory inference model. For each dataset, we inferred trajectories and pseudotime values using Slingshot^38^, and embedded the single-cells using PHATE^39^, a method that preserves local and global relationships, and continuous progressions in high-dimensional data. We excluded two clusters (“unknown 1” and “unknown 2”) from aRMS primary cultures, as these subpopulations were present only in two specific samples, which might be the result of incomplete batch effect removal (**Extended Data Figure 3a**). In primary cultures derived from both RMS subtypes, the predicted trajectory revealed a branched progression from *MuSC-like* cells to either actively *cycling progenitors* or to *differentiated* cells (**Figure 3a**), resembling the branched trajectory described in two recent scRNAseq studies on regenerating skeletal muscle cells^30, 31^.

In the aggressive aRMS subtype no hierarchical organization has yet been described, therefore we aimed at understanding the relationship between these cancer cells and myogenic cell types. We re-analyzed a published scRNAseq dataset of regenerating skeletal muscles^31^ and projected aRMS cancer cells from five primary cultures onto this roadmap (**Figure 3b**). Myogenic and tumor cells overlapped on the branched trajectory (**Figure 3c, Extended Data Figure 3b**). Notably, the *cycling progenitor* population, which was only transiently detected in skeletal muscle cells upon injury (**Extended Data Figure 3c**), represents a prominent and stable population in aRMS tumors.

**Figure 3:**
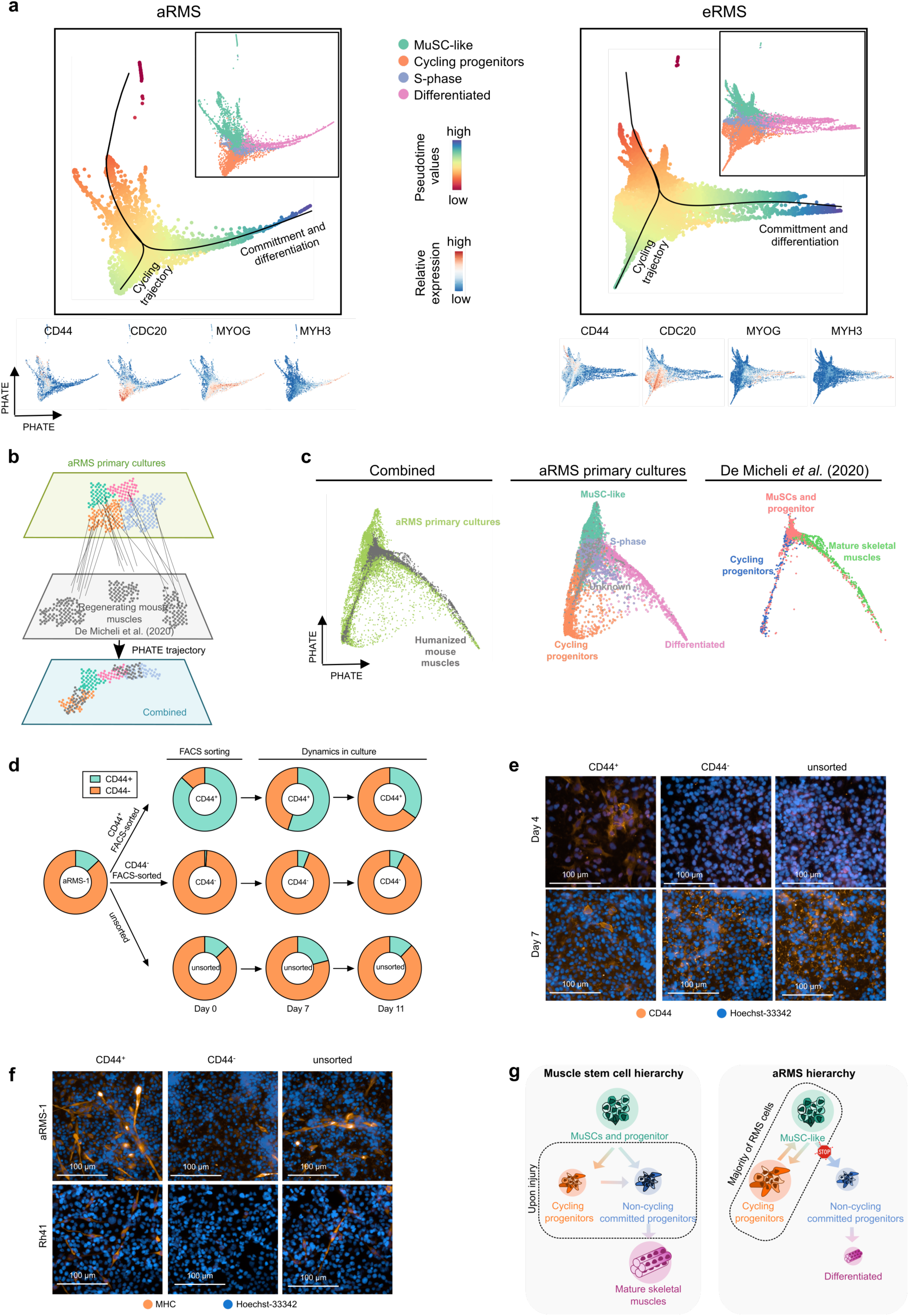
RMS primary cultures recapitulate a branched myogenic trajectory. **a.** ScRNAseq data generated from *n* = 5 aRMS (left) or *n* = 9 eRMS (right) primary cultures were integrated using SCT-correction. All clusters that were commonly shared across all the aRMS patients were selected and re-analyzed by PHATE dimensionality reduction (*t* = 30, *knn* = 20). Black lines represent pseudotime trajectories calculated using the *Slingshot* package (starting cluster set as *Progenitor*); dashed arrows represent the trajectory direction as predicted by *Slingshot*. Cells are colored based on the identified Louvain clusters or based on pseudotime values calculated by *Slingshot*. The expression of genes delineating a myogenic progression (CD44, *MuSC-like*; CDC20, *cycling progenitors*; MYOG, committed progenitors; MYH3, differentiated cells) is also shown (lower panel). **b.** Computational workflow. ScRNAseq data generated from mouse muscle tissues (De Micheli *et al*., 2020) were subset on myogenic populations. To compare expression across mouse and human species, the gene names in De Micheli *et al.* (2020) were humanized and integrated with scRNAseq data generated from aRMS primary cultures. The combined dataset was then reduced and visualized by PHATE dimensionality reduction. **c.** PHATE dimensionality reduction (*t* = 30, knn = 20) plots of the single-cell datasets derived from muscle tissues (De Micheli et al. *Cell Rep.* 2020), aRMS primary cultures or from their combined integration. ARMS cells are color-coded based on Louvain clusters; muscle cells are colored based on the clusters identified in the original publication. **d.** Flow cytometry analysis of FACS-sorted CD44^+^ and CD44^-^ subpopulations in aRMS-1 cells. Unsorted reference is also shown. Data are represented as mean of *n* ≥ 2 biological replicates. **e.** Immunofluorescence analysis of CD44 expression four and seven days after sorting CD44^+^ and CD44^-^ subpopulations in aRMS-1 cells. **f.** Immunofluorescence analysis of MyHC expression seven days after sorting of CD44^+^ and CD44^-^ subpopulations in aRMS-1 cells. **g.** Proposed model of aRMS hierarchical structure compared to the muscle stem cell hierarchy proposed in De Micheli *et al.* (2020). When muscle stem cells (MuSCs) get activated, for example upon injury, they expand as cycling progenitors and finally differentiate into mature skeletal muscles. The majority of aRMS cells are blocked in a continuous loop between *MuSC-like* and *cycling progenitors*, except for a minority of cells that escapes the differentiation block, becomes committed and differentiates.

To better characterize the kinetics of the cell trajectory in aRMS, we took advantage of the cell surface marker CD44, which we found to be predominantly expressed on *MuSC-like* RMS cells in our scRNAseq and CyTOF analysis. We first measured CD44 expression across RMS primary cultures and cell lines and then focused our study on the primary cultures labeled aRMS-1 and aRMS-3, because they exhibited two distinct subpopulations distinguished by CD44 expression, CD44^+^ and CD44^-^ cells (**Extended Data Figure 3d**). We assessed the feasibility of using CD44 as a marker of *MuSC-like* cells by measuring the transcript levels of key myogenic markers in CD44^+^ and CD44^-^ cell populations that were purified by Fluorescence-Activated Cell Sorting (FACS). CD44^+^ cells exhibited increased expression of CD44 and PAX7 (log_2_ FC: 3.1±0.5 in aRMS-1, 2.4±0.3 in aRMS-3), and decreased expression of MYOD1 (log_2_ FC: -0.5±0.3 in aRMS-1, -0.7±0.1 in aRMS-3) and MYOG (log_2_ FC: -1.1±0.3 in aRMS-1, -2.0±0.1 in aRMS-3), compared to CD44^-^ cells (**Extended Data Figure 3e**), supporting the rationale of using CD44 as a *MuSC-like* marker. We sought to determine whether the trajectory of aRMS primary cultures was unidirectional or whether cycling progenitors could de-differentiate to a *MuSC-like* state. To address this question, we sorted *MuSC-like* aRMS cells (CD44^+^) from the other subpopulations (CD44^-^), which mainly encompass *cycling progenitors*, and monitored their behavior in culture. We quantified CD44 expression at the protein level in the sorted populations over time, using both flow cytometry and immunofluorescence, and found that both CD44^+^ and CD44^-^ subpopulations re-gained their initial mixed composition after 2 weeks in culture (**Figure 3d, Figure 3e, Extended Data Figure 3f**). Since CD44^+^ cells lost CD44 expression faster than CD44^-^ gained it, we wondered whether the presence of CD44^+^ cells in the CD44^-^ sorted subpopulation was due to self-renewal of some remaining *MuSC-like* cells that were not completely removed during sorting. However, we observed similar growth rates (data not shown) and cell cycle distributions of both subpopulations (**Extended Data Figure 3g**). Although we cannot rule out *in vitro* selection of a less differentiated phenotype, our data are also consistent with the hypothesis of de-differentiation wherein *cycling progenitors* can regain a *MuSC-like* phenotype. Finally, to determine whether *cycling progenitors* can directly transition towards committed and differentiated states, we FACS-sorted CD44^+^ and CD44^-^ subpopulations and stained them for MyHC seven days after sorting. We observed differentiated MyHC^+^ cells predominantly in CD44^+^ and unsorted cells, but not in CD44^-^ cells (**Figure 3f**). These findings suggest that, in aRMS, *cycling progenitors* lack the ability to directly transition towards *differentiated states*, which is characteristic of *MuSC-like* cells.

In summary, our data show that RMS are hierarchically organized and that aRMS contain two plastic oncogenic subpopulations, namely the transcriptionally immature *MuSC-like* subpopulation and the *cycling progenitor* subpopulation (**Figure 3g**).

### PAX3:FOXO1 sustains aRMS cells in the cycling program and its depletion leads to *MuSC-like* and *differentiated* cells

PAX3:FOXO1 is the major driver of aRMS, known to repress myogenic differentiation. To determine how PAX3:FOXO1 regulates the subpopulations identified in aRMS, we used two aRMS cell lines (Rh4 and KFR) expressing a doxycycline (DOX)-inducible short hairpin RNA directed against PAX3:FOXO1 (shP3F1) or against a control short hairpin RNA (shSCR)^40^. We decreased PAX3:FOXO1 levels by treatment with DOX for 48 hrs and measured the transcriptome of 4,582 Rh4 and 6,005 KFR cells using droplet-based single-cell RNA sequencing (**Figure 4a**). Upon knockdown (KD) of the fusion protein (**Figure 4b and Extended Data Figure 4a**), both Rh4 and KFR cells underwent consistent transcriptional changes at the global level, including upregulation of the skeletal muscle genes MYL1, MYOG and DES, and repression of the known PAX3:FOXO1 downstream targets STATH, PIPOX and ASS1 in Rh4 cells (**Extended Data Figure 4b and 4c; Supplementary Table 2**). At the single-cell level, the KD of the fusion protein induced clear cellular shifts (**Figure 4c**). The majority of the cells acquired a signature characteristic of *differentiated* cells, and a minority of the cells acquired a *MuSC-like* signature; the *cycling progenitor* signature was completely lost (**Figure 4d**). In KFR, the impact of the KD on the *differentiation*, *MuSC-like* and *cycling progenitor* programs was observed both at the single-cell and bulk level, while in Rh4 the emergence of the *MuSC-like* cluster in Rh4 and the reduction in *cycling progenitor* programs was less evident at the bulk level (**Figure 4e**). Collectively, these findings support a model where PAX3:FOXO1 sustains aRMS cells in an intermediate *cycling progenitor* state (**Figure 4f**). Upon PAX3:FOXO1 KD, cells either undergo differentiation, or halt in a *MuSC-like* state, a finding that can inform the design or implementation of PAX3:FOXO1-targeted therapy.

**Figure 4:**
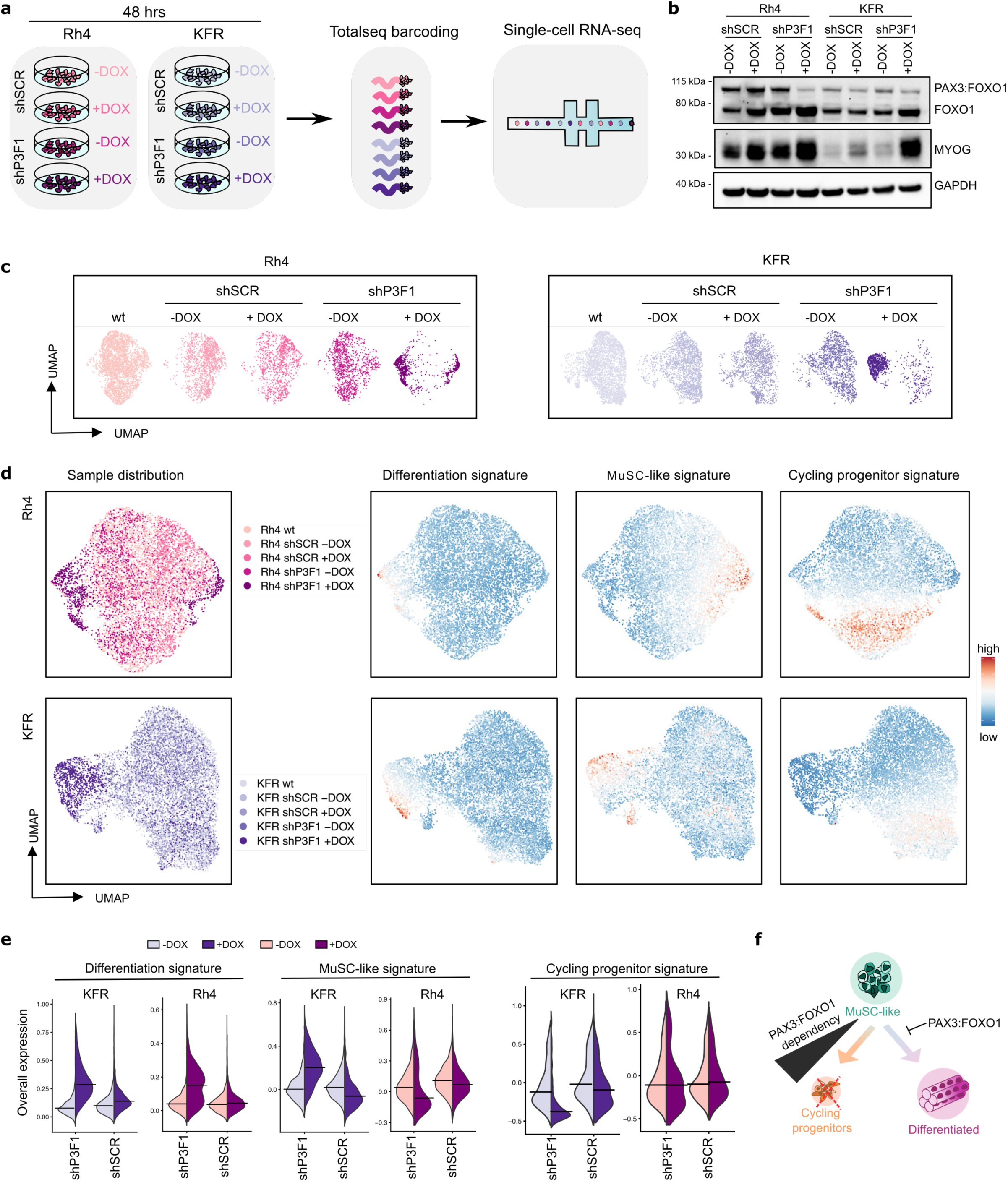
Single-cell responses upon PAX3:FOXO1 downregulation in aRMS cell lines. **a.** Schematic workflow. Rh4 and KFR cells were transduced with a DOX-inducible shRNA against P3F1 (shP3F1) or against a scramble control (shSCR). After treatment with DOX for 48 hrs, cells were collected, stained with Totalseq hashtags and pooled together before sequencing on a 10X Chromium instrument. **b.** Representative western blot of Rh4 and KFR cells cultured with or without DOX for 48 hrs. **c.** UMAP plot of 8,093 Rh4 and 9,501 KFR cells profiled by scRNAseq from shSCR and shP3F1 lines upon treatment with DOX and combined with scRNAseq data generated from *wild type* lines. **d.** UMAP plot of the integrated Rh4 and KFR cells, colored by: the genetic perturbation and DOX treatment (left panel) or the overall expression (color scale) of differentiation, *MuSC-like*, and *cycling progenitor* signatures (right panel). The expression of each signature was calculated based on the genes identified in the combined aRMS cell line dataset. **e.** Distribution of overall expression scores for each signature in control (shSCR) and PAX3:FOXO1 KD (shP3F1) lines upon treatment with DOX. **f.** Proposed model of P3F1 heterogeneity across aRMS cell lines. Upon P3F1 removal, the differentiation block is released and the oncogenic loop disrupted. The *cycling progenitor* subpopulation disappears and the remaining cells display *MuSC-like* or *differentiation* features.

### Image-based high content screening reveals that chemotherapy enriches aRMS tumors for the plastic *MuSC-like* subpopulation

Our single-cell analysis showed that aRMS primary cultures contain *MuSC-like* cells that either commit to *differentiation* or give rise to actively *cycling progenitors*. To understand how we could target these cellular transitions we directed our efforts toward the identification of (i) drug-induced beneficial effects, which are compounds promoting cellular differentiation, (ii) drug-induced detrimental effects, which we defined as compounds that de-differentiate aRMS, enriching the tumors into for a plastic *MuSC-like* subpopulation. To quantify the aRMS cellular states in a high-throughput manner, we first established MYOscopy, a high content microscopy screening platform aimed to detect the expression of Ki-67, myogenin and MyHC, markers of proliferating, committed and differentiated myogenic cells, respectively. This approach was able to distinguish *quiescent MuSC-like* (myogenin^-^/Ki-67^-^), *cycling MuSC-like* (myogenin^-^/Ki-67^+^), *cycling progenitors* (myogenin^+^/Ki-67^+^/MyHC^-^)*, committed progenitors* (myogenin^+^/Ki-67^-^/MyHC^-^) and *differentiated* (MyHC^+^) cellular states in aRMS primary cultures (**Extended Data Figure 5a and 5b**), demonstrating its potential to serve as a phenotypic screening platform.

To identify drug-induced transitions, we exposed two aRMS primary cultures to a drug library of 244 compounds in a dose range of 10 nM-10 μM and used MYOscopy to quantify the cell composition after 72 hrs of treatment (**Figure 5a; Supplementary Table 3**). We identified 15 compounds that consistently enriched both aRMS primary cultures for *MuSC-like* subpopulations (**Figure 5b**). Notably, the commonly used chemotherapeutics vinblastine sulfate, daunorubicin and doxorubicin were among these 15 drugs (**Figure 5c**), and they all exhibited dose-dependent effects (**Extended Data Figure 5c**).

**Figure 5:**
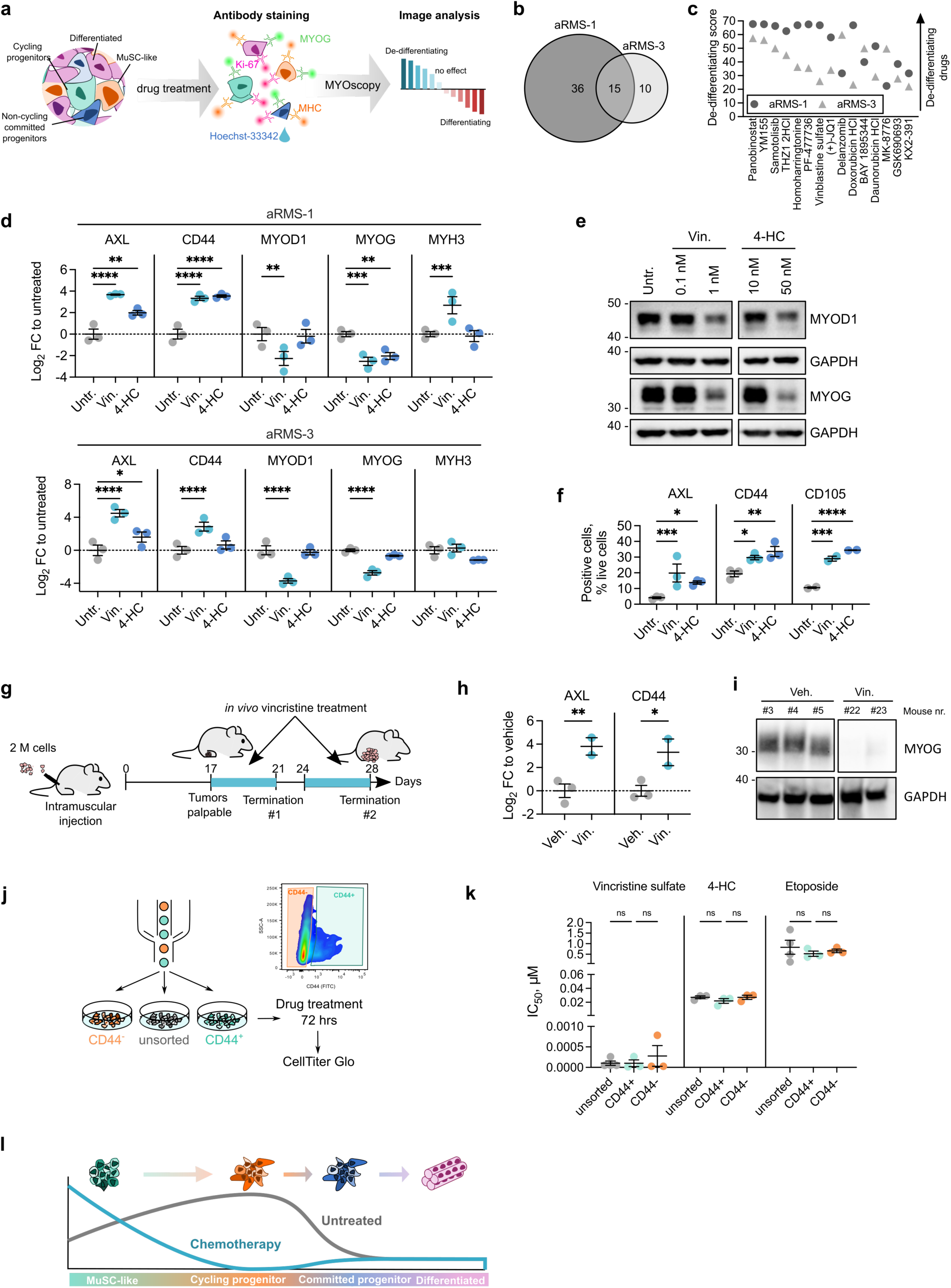
Clinically used chemotherapeutics induce a transition towards progenitor cell states in aRMS. **a.** Single-cell high-content image experimental workflow. **b.** Number of de-differentiating drug hits in aRMS-1 and aRMS-3 cells at 1 μM. **c.** Shared de-differentiating drugs at 1 μM ranked based on the average de-differentiating score (see methods) across aRMS-1 and aRMS-3 cells. A score of zero represents the baseline score of untreated controls. **d.** QRT-PCR data generated with aRMS-3 and aRMS-1 cells exposed to the chemotherapeutics vincristine sulfate (10 nM in aRMS-1, 1 μM in aRMS-3 cells) or 4-HC (50 nM in both aRMS-1 and aRMS-3 cells) for 48 hrs. Data are represented as mean ± SEM of *n* = 3 biological replicates; ordinary two-way ANOVA with uncorrected Fisher’s LSD. **e.** Western blot analysis of MyoD1 and myogenin protein levels in aRMS-1 cells exposed to the chemotherapeutics vincristine or 4-HC for 48 hrs at the indicated concentrations. GAPDH was used as loading control; samples were loaded on the same gel. **f.** FACS analysis of aRMS-1 cells treated with 1 nM vincristine or 10 nM 4-HC for 48 hrs. Dead cells were excluded by Zombie NIR staining, and AXL^+^, CD44^+^ and CD105^+^ cells were quantified based on isotype controls. Data are represented as mean ± SEM of *n* = 3 biological replicates (*n* = 2 for CD105); ordinary two-way ANOVA with uncorrected Fisher’s LSD. **g.** Schematic of *in vivo* vincristine sulfate treatment. *N* = 2 mice were treated with 10 mg/kg drug for either one or two weeks. Tumors were collected and the mRNA expression of myogenic markers was analyzed by qRT-PCR and compared to vehicle-treated animals. **h.** QRT-PCR data generated with aRMS-1 tumors collected after one or two weeks of *in vivo* vincristine sulfate or vehicle treatment. Data are represented as mean ± SEM of the indicated number of mice; ordinary two-way ANOVA with uncorrected Fisher’s LSD. **i.** Western blot analysis of myogenin protein levels in aRMS-1 tumors treated *in vivo* with vincristine sulfate or with vehicle control. Samples were loaded on the same gel. **j.** Experimental workflow. aRMS-1 cells were first FACS-sorted based on expression of CD44, then exposed to the indicated drugs at five different concentrations. After 72 hrs, viable cells were quantified by CellTiter Glo. Unsorted cells were used as a reference. **k.** Half maximal inhibitory concentration (IC_50_) values of vincristine, 4-HC and etoposide in aRMS-1 cells calculated based on dose-response curves. Data are represented as mean ± SEM of n ≥ 3 biological replicates. **i.** Proposed model of selection following treatment with chemotherapy in aRMS cells. *, *P* < 0.05; **, *P* < 0.01; ***, *P* < 0.001; ****, *P* ≤ 0.0001 *Untr*.: untreated; *Vin*: vincristine sulfate; *4-HC*.: 4-hydroperoxycyclophosphamide; *Veh.*: vehicle; *FC*: fold change; *hrs*: hours.

As resistance to chemotherapy is a major concern in aRMS, we characterized the effect of clinically-used chemotherapeutics, such as vincristine, etoposide and cyclophosphamide, on the cellular composition of aRMS primary cultures in more detail. After a 72 hrs exposure to vincristine or etoposide, we observed a clear decrease in the number of nuclei, a significant increase in the proportion of *MuSC-like* cells, and a corresponding decrease in the proportion of *cycling progenitors*; there was no significant difference in the composition of *non-cycling committed progenitors* and *differentiated* cells (**Extended Data Figure 5d and 5e**). To further validate the results at the transcriptome level, we exposed two aRMS primary cultures to vincristine and 4-Hydroperoxycyclophosphamide (4-HC), the active metabolite of cyclophosphamide and evaluated their effects on the mRNA level of selected markers. Both drugs increased AXL (log_2_ FC: 3.6±0.1 after vincristine, 2.0±0.2 after 4-HC) and CD44 mRNA levels (log_2_ FC: 3.3±0.2 after vincristine, 3.5±0.1 after 4-HC), markers of the *MuSC-like* subpopulation, and decreased MYOG (log_2_ FC: -2.5±0.4 after vincristine, -2.0±0.3 after 4-HC), characteristic for the committed/differentiated subpopulation. Vincristine slightly decreased the expression of the activated myogenic marker MYOD1, whereas we found no consistent change in the differentiation marker MYH3 (**Figure 5d**). We confirmed drug-induced downregulation of MYOD1 and MYOG (**Figure 5e, Extended Data Figure 5f and 5g**) at the protein level by western blot analysis and upregulation of the progenitor markers CD44 (1.5-fold after vincristine, 1.7-fold after 4-HC), AXL (4.8-fold after vincristine, 3.3-fold after 4-HC) and CD105 (2.7-fold after vincristine, 3.3-fold after 4-HC) by flow cytometry analysis (**Figure 5f**). To determine the effect of vincristine treatment *in vivo*, we injected aRMS-1 cells orthotopically into immunodeficient NSG mice and collected the tumors after one and two weeks after vincristine treatment (**Figure 5g**). Indeed, *in vivo* treated tumors showed increased mRNA expression of the *MuSC-like* markers CD44 and AXL mRNA (**Figure 5h**) and decreased protein expression of MYOG (**Figure 5i**). Finally, we wished to determine whether progenitor states are intrinsically more resistant to chemotherapy. We assessed the *in vitro* efficacy of vincristine or 4-HC on sorted CD44^+^ and CD44^-^ cells (**Figure 5j**). Strikingly, all drugs were equally effective on both CD44^+^ and CD44^-^ subpopulations, with no significant difference in their IC_50_ values (**Figure 5k**, **Extended Data Figure 5h**). These results suggest that chemotherapy enriches aRMS cells for a *MuSC-like* state (**Figure 5l**) by rewiring the cellular trajectory and not necessarily by selecting intrinsically chemo-resistant subpopulations.

### Trametinib overcomes the differentiation block in aRMS primary cultures

Differentiation therapy has been proposed as a promising therapeutic strategy in cancers with dysregulated development^23, 24^. To identify compounds that induced differentiation of aRMS, we re-analyzed our drug screening performed on two aRMS primary cultures (**Figure 5a**). Specifically, we focused on compounds that were able to increase the *non-cycling committed progenitors* and *differentiated* subpopulations, while simultaneously decreasing *cycling progenitors*. Of the 244 FDA-approved anticancer agents tested, we identified 72 compounds that were able to promote differentiation; 13 of the 72 compounds (18%) were effective on both aRMS-1 and aRMS-3 cells (**Figure 6a**). These compounds included the MEK inhibitors trametinib and cobimetinib, the FGFR inhibitor erdafitinib, the VEGFR/FGFR inhibitors ponatinib and lenvatinib mesylate and the aurora kinase inhibitors alisertib and AT9283 among others (**Figure 6b**). Trametinib, cobimetinib and erdafitinib appeared as top-differentiating compounds, and all induced morphologically robust myogenic differentiation in a concentration-dependent manner (**Figure 6c; Extended Data Figure 6a and 6b**). Notably, in contrast with our finding using aRMS primary cultures, a recent report failed to detect differentiation in aRMS cell lines treated with trametinib^26^. We therefore compared the therapeutic potential of trametinib-induced differentiation in seven aRMS primary cultures and two aRMS cell lines. Trametinib treatment only slightly reduced the number of nuclei in both primary cultures and cell lines, but it consistently shifted the primary cultures towards *non-cycling committed progenitor* and *differentiated* states, while reducing the *cycling progenitor* state, an effect that was observed to a lower extent in the cell lines (**Figure 6d**, **Extended Figure 6c, 6d**). Upon treatment with trametinib, we found a consistent increase in myogenin expression across both primary cultures and cell lines, while the increase in MyHC expression was limited to the primary cultures (**Figure 6e**, **Figure 6f**). Notably, the protein expression levels of both the *MuSC-like* marker Pax-7, when present, and of PAX3:FOXO1 decreased upon trametinib treatment. In primary cultures, mRNA expression of the *MuSC-like* markers CD44 and AXL decreased, while the mRNA expression of MYOG and MYH3 mRNA increased, consistent with cellular shifts towards differentiated states (**Figure 6g**). Cell cycle analysis confirmed that trametinib treatment led to an arrest in G_0_/G_1_ phase in both primary cultures and cell lines, as expected from cells undergoing terminal myogenic differentiation (**Figure 6h and Extended Figure 6e**). To determine whether trametinib induces differentiation by on-target effects, we measured the effect of trametinib on ERK phosphorylation. In all tested primary cultures and cell lines we observed a decrease in phosphorylated ERK upon treatment with low doses of trametinib (in the nM range) (**Figure 6i**), which correlated with the observed trametinib-induced differentiation (*R*^2^ = 0.83; *P* = 0.03; *n* = 5) (**Figure 6j**). These findings underscore the advantage of using primary cells rather than cell lines for drug screening and suggest that the RAS pathway might be a target for a differentiation therapy approach in aRMS.

**Figure 6:**
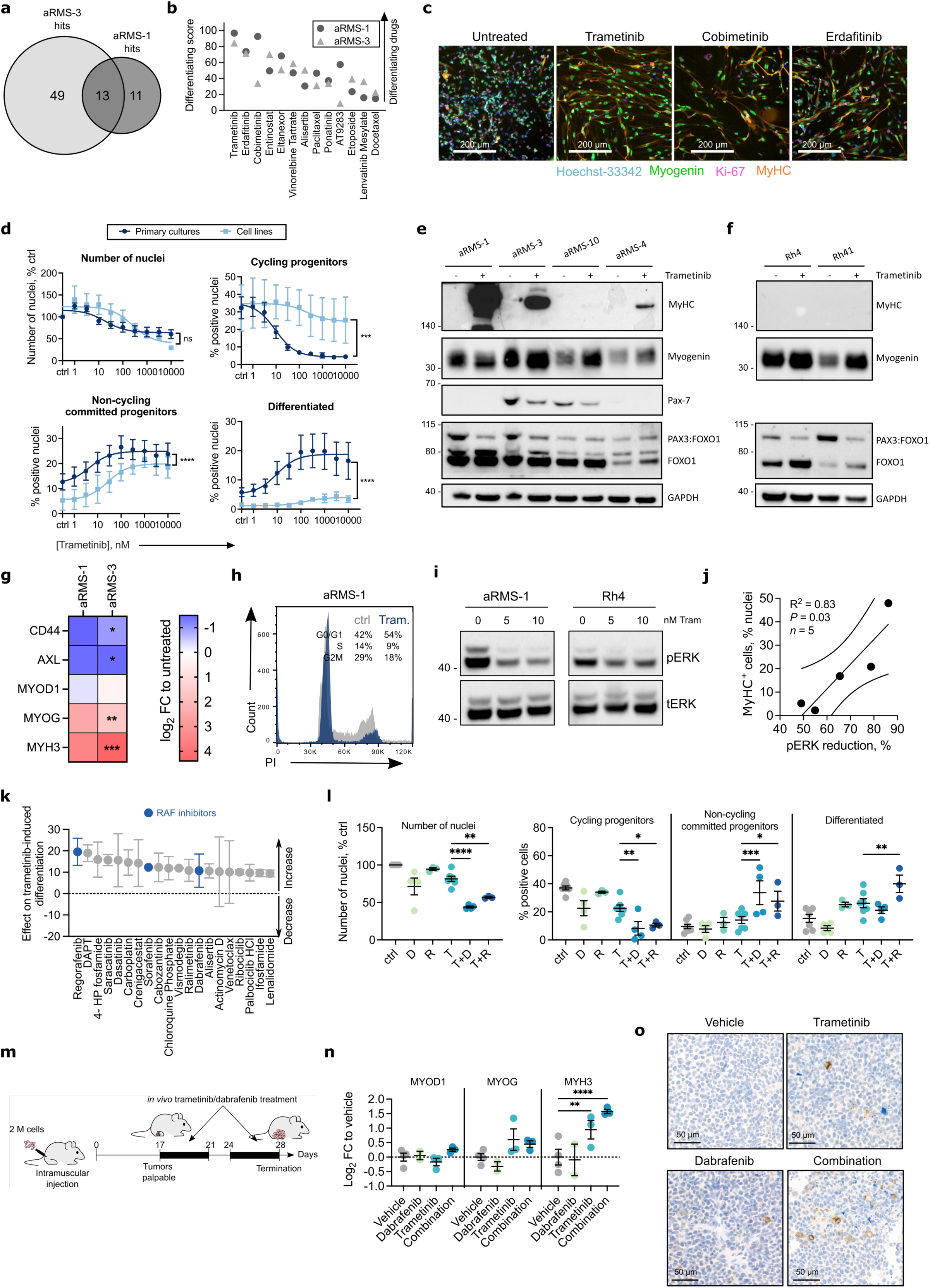
Trametinib induces myogenic differentiation and cell cycle arrest in aRMS primary cultures. **a.** Number of drug hits promoting differentiation in aRMS-1 and aRMS-3 at 1 μM. **b.** Shared differentiating drugs at 1 μM ranked based on the average differentiating score (see Methods) across aRMS-1 and aRMS-3 cells. A score of zero represents the baseline score of untreated controls. **c.** Representative IF images of aRMS-1 cells following 72 hrs drug treatment at 1 μM. **d.** Dose-response curves of the composition of aRMS primary cultures and cell lines exposed to the MEK inhibitor Trametinib for 72 hrs. The number of nuclei, percentage of *cycling*, *non-cycling committed progenitors* and *differentiated* cells was measured with MYOscopy. Data are represented as mean ± SEM of *n* = 7 primary cultures and *n* = 2 cell lines; paired *t*-tests. **e and f.** Western blots of aRMS primary cultures (**e**) and cell lines (**f**) exposed to 50 nM Trametinib for 96 hrs. **g.** QRT-PCR data generated with aRMS-1 or aRMS-3 cells exposed to 50 nM Trametinib for 96 hrs. Data are represented as mean ± SEM of *n* ≥ 2 biological replicates; multiple unpaired *t*-tests. **h.** Representative cell cycle analysis FACS plot of aRMS-1 cells exposed to 25 nM trametinib for 96 hrs and stained with propidium iodide. **i.** Western blot analysis of phosphorylation of ERK in a representative aRMS primary culture (aRMS-1) and cell line (Rh4) after exposure to 5 or 10 nM Trametinib for 3 hrs. **j.** Correlation between the reduction in ERK phosphorylation after 3 hrs exposure to 10 nM Trametinib and the percentage of differentiated cells (MyHC^+^) after exposure to 1 μM Trametinib for 72 hrs measured by MYOscopy. Data points, representing different aRMS primary cultures or cell lines, are interpolated with a linear regression. Correlation coefficient (R^2^), statistical significance (*P*) and number of data points (*n*) are indicated. **k.** Top 20 trametinib-potentiating drugs ranked based on their effect on trametinib-induced differentiation (see Methods). Each drug was tested in combination with 50 nM trametinib in aRMS-1 cells. A score of zero represents the baseline score of trametinib alone; drugs with a positive score potentiate trametinib-induced differentiation, whereas drugs with a negative score diminish differentiation. Data points are represented as mean ± SEM of *n* = 2 biological replicates. **l.** Quantification of IF images of aRMS-1 cells exposed to control (ctrl), 10 μM dabrafenib (D), 1 μM regorafenib (R), 10 nM trametinib (T), or to the indicated combinations for 72 hrs. **m.** Schematic of *in vivo* trametinib/dabrafenib treatment. Mice were treated with 1 mg/kg trametinib, 15 mg/kg dabrafenib or with their combination. After two weeks of drug treatment (day 28 post injection), tumors were collected and myogenic markers were analyzed by qRT-PCR and immunohistochemistry. **n.** QRT-PCR analysis of aRMS-1 tumors. Data are represented as mean ± SEM of the indicated number of mice; ordinary two-way ANOVA with uncorrected Fisher’s LSD. **o.** Expression of MyHC as determined by immunohistochemistry in aRMS-1 PDX tumors following *in vivo* treatment with 1 mg/kg trametinib, 15 mg/kg dabrafenib or with their combination. *, *P* < 0.05; **, *P* < 0.01; ***, *P* < 0.001; ****, *P* ≤ 0.0001

Since clinical trials with single targeted agent treatments have been largely unsuccessful, we next sought to identify trametinib-potentiating agents by performing a combinatorial drug screening in aRMS-1 cells. Specifically, we used a library of 243 compounds either alone or in combination with 50 nM trametinib and measured differentiation using MYOscopy (**Supplementary Table 4**). We found 97 drugs that potentiated trametinib-induced differentiation; of the top 20 hits, we identified several RAF inhibitors (**Figure 6k**), including regorafenib, currently in clinical trial in RMS patients, and dabrafenib, already used in combination with trametinib in patients with melanoma^41^ and non-small cell lung cancer^42^ patients. We measured the differentiating effect of trametinib-dabrafenib and trametinib-regorafenib combinations at different drug ratios and observed a synergistic score when 10 nM trametinib was combined with 1 µM regorafenib or 10 µM dabrafenib (**Extended Data Figure 6f**). At this drug ratio, both combinations significantly decreased the number of nuclei and of *cycling progenitors* and shifted primary cells towards *non-cycling committed progenitor* and *differentiated* stages compared to single-treatment with trametinib (**Figure 6l and Extended Data Figure 6g**). To examine the *in vivo* relevance of trametinib-dabrafenib combination, we injected aRMS-1 cells orthotopically and collected the tumors after two weeks of trametinib and dabrafenib treatment, either as single agents or in combination (**Figure 6m**). Tumors that were exposed to the drug combination showed significantly higher mRNA levels of the differentiation marker MYH3 (**Figure 6n**). Immunohistochemistry analysis also showed an increase in the proportion of myogenin^+^, MyHC^+^ and a decrease in the proportion of Ki-67^+^ cells in tumors exposed to the drug combination (**Figure 6o, Extended Data Figure 6h**), indicative of myogenic differentiation. Overall, these data suggest that trametinib-induced suppression of the RAS pathway in combination with RAF inhibitors is a potential novel treatment strategy for aRMS that targets cellular differentiation.

## DISCUSSION

Single-cell RNA and protein profiling are powerful methods for investigating developmental heterogeneity and plasticity in human cancers. Here, we leveraged the use of scRNAseq and CyTOF analysis on clinically relevant models of RMS to dissect the cellular, molecular and functional intra-tumor heterogeneity. Our findings provide robust evidence in support of the hypothesis that both aRMS and eRMS tumors harbor an abundance of developmentally halted cells with myogenic potential that fail to differentiate. Most importantly, we identify a myogenin^+^ *cycling progenitor* cell state that is only transiently found in myogenesis during muscle development and regeneration^31^ and that distinguishes a more oncogenic (aRMS) from a less oncogenic (eRMS) tumor phenotype. Myogenin is typically a hallmark transcription factor of a non-proliferative myogenic commitment state that drives expression of differentiation markers, such as myosin heavy chains. In aRMS, this *cycling progenitor* state continues to proliferate, even though it has initiated myogenic differentiation. In addition, we identify two previously unrecognized RMS cell subpopulations, one resembling early muscle stem cells (*MuSC-like*) and one resembling differentiated muscle cells. While these cellular states were clearly present in RMS primary cultures, their abundance was notably decreased in RMS cell lines, providing support for the preclinical use of primary cultures in lieu of cell lines. An emerging body of evidence suggests that childhood tumor types contain developmental hierarchies^4–7^. Here we provide evidence that this phenomenon is a characteristic of RMS. Moreover, a characterization of the developmental hierarchies in the aggressive aRMS models reveals a distinction of relevance to clinical outcome. When *MuSC-like* cells become aberrant *cycling progenitors*, they correlate with worst patient prognosis; on the other hand, if they are able to commit to a *non-cycling committed/differentiated* cell state, they drive better patient prognosis. Our data supports a model for aRMS in which *cycling progenitors* can revert back to a *MuSC-like* state, fueling a continuous oncogenic loop. We therefore propose an aRMS tumor model in which there is no “cancer stem cell” population, but instead both *MuSC-like* and *cycling progenitor* cells co-exist in plastic oncogenic states that must be therapeutically targeted, a feature recently described as a newly recognized hallmark of cancer^2^. Hence, for aRMS, a differentiation-inducing approach that overcomes the block and re-wires the trajectory is likely to be effective.

Our results demonstrate that following genetic perturbation of the fusion protein PAX3:FOXO1, cells unexpectedly revert back to a *MuSC-like* state, or undergo myogenic differentiation, as previously described^34, 43^. These findings suggest that PAX3:FOXO1 maintains aRMS tumor cells in the *MuSC-like/cycling progenitor* trajectory loop. The other cellular states may have intrinsically lower PAX3:FOXO1 expression or may be less dependent on expression of the fusion protein itself. However, it still remains to be determined whether a population of cells that is completely independent of PAX3:FOXO1 exists and whether it has the potential to proliferate, as suggested based on data from ectopic models^22^. Interestingly, both mesenchymal stem cells^10^ and differentiated myogenic cells^12^ have previously been described as possible cells of origin for aRMS, a finding supported by our studies.

The image-based screening approach we developed here for aRMS can serve as a guide for other cancers to better understand the mechanisms determining cancer cell fate decisions. Patients with aRMS are known to respond to the first treatment, but they often relapse^16^, suggesting that therapy fails to eliminate all tumor cells. It is therefore not surprising that, as found here, clinically used chemotherapeutics cause enrichment in aRMS tumors of *MuSC-like* cells, as those cells are primarily low-cycling and therefore not targeted by chemotherapy. Based on our data, it is this state that can expand and recapitulate the trajectories that reflect the tumor complexity. Unlike other cancers, in which the progenitor/stem cell population has been regarded as resistant to treatment, our findings suggest that *MuSC-like* aRMS cells are not intrinsically chemo-resistant. We therefore propose a model in which chemotherapy rewires the cellular hierarchy by either selecting for *MuSC-like* cells or de-differentiating *cycling* myogenin^+^ *progenitors*, creating a population that leads to relapse.

The presence of differentiated cells in RMS tumors under basal conditions indicates that some cells can escape the differentiation block and that aRMS tumors may be amenable to differentiation therapy. To determine if we could pharmacologically capitalize on this process, we first screened for compounds that promote differentiation. We identified trametinib-induced MEK-inhibition as a strategy to rewire the tumor hierarchy in aRMS primary cell cultures, in a manner not seen with aRMS cell lines by us or others^26^. This leads us to conclude that cell lines may be less prone to surmount the differentiation block and therefore lack utility for drug screening. Our findings underscore the use of primary cells from tumors as models to study RMS tumorigenesis. Our combination screen identified RAF inhibitors as top trametinib-potentiating compounds in aRMS, in line with recent studies^25^ showing that vertical inhibition of the RAF-MEK-ERK cascade effectively induces differentiation in eRMS models. We were able to confirm our *in vitro* findings by *in vivo* induction of RMS differentiation following a combination of RAF and MEK targeting by trametinib and dabrafenib treatment, however, its therapeutic relevance remains to be established. Clinical studies with the RAF inhibitor regorafenib are currently ongoing in RMS (NCT01900743), while studies with the MEK inhibitor cobimetinib are underway in eRMS patients (NCT02639546). Based on our findings, we expect that RAF inhibitors, such as regorafenib, would benefit from a combination with MEK inhibitors, an approach that could lead to increased aRMS cellular differentiation. Notably, the use in the clinic of differentiation therapy for aRMS and its combination with chemotherapy have to be carefully evaluated, as our work suggests that they induce opposite lineage shifts. To be successful, one would first need to eliminate the rapidly dividing *cycling progenitors* with chemotherapy, and only then “mobilize” the remaining *MuSC-like* cells in order to cause their exhaustion using differentiating agents.

In summary, our work provides the first comprehensive single-cell transcriptomic and proteomic atlas of RMS. This atlas resolves the cellular and functional heterogeneity of RMS and reveals key cellular and molecular signatures, with therapeutic implications for chemoresistance and tumor relapse. Our study is the first of such a kind to shed light on the cellular fate mechanisms underlying impaired differentiation in the aggressive alveolar RMS subtype.

## METHODS

### PDX generation and dissociation

The PDXs used in this study were generated from patient biopsy samples collected at St Jude Children’s Research Hospital, University Children’s Hospital Zurich, Emma Children’s Hospital Amsterdam, Institut Curie Paris and Charité University Hospital Berlin as described previously.^44^ Patient characteristics and information on the clinical status can be found in **Extended Data Table 1**. Patient biopsies were first expanded in immunodeficient NOD scid gamma (NSG), Janvier Rj:NMRI-*Foxn1^nu/nu^* or *Taconic* NOD.Cg-*Prkdc^scid^ Il2rg^tm1Sug^*/JicTac mice as previously described. Tumors were isolated from mice when reaching a size of 700-1300 mm^3^, mechanically minced into smaller pieces using scalpels and re-transplanted in secondary recipient mice. To generate single cell suspensions, PDX tumors were mechanically and enzymatically digested using 200 μg/ml liberase DH (Roche, 5401054001) and 1 mM MgCl_2_ in 1x HBSS buffer (Sigma, H6648) for 30–60 min at 37 °C. Cell suspension was filtered through a 70μm cell strainer, to remove remaining tumor pieces, washed with PBS, and used immediately to produce cultured cells or frozen in freezing medium CryoStor CS10 (StemCell, #07930).

### Primary RMS culture

To produce primary cultures, dissociated PDX tumors were grown on plates coated with Matrigel (Corning, 354234) diluted 1:10 in Advanced DMEM/F-12 medium (Thermofisher Scientific, 12634010) and left at room temperature (RT) for 30-60 min to solidify. Cells were cultured in Advanced DMEM/F-12 (Thermofisher Scientific, 12634010) medium supplemented with 100 U/ml penicillin/streptomycin (Thermofisher, 15140122), 2 mM Glutamax (Life technologies, 335050-061), 0.75x B-27 (Thermofisher Scientific, 17504044), 20 ng/ml bFGF (PeproTech, AF-100-18B) and 20 ng/ml EGF (PeproTech, AF-100-15,) (“Complete F12” medium) or in Neurobasal medium (Thermo Fisher Scientific) supplemented with 100 U/ml penicillin/streptomycin (Thermofisher, 15140122), 2 mM Glutamax (Life technologies, 335050-061), 2x B-27 (Thermofisher Scientific, 17504044) (“Complete NB” medium). In some cases, complete NB medium was supplemented with 20 ng/ml bFGF bFGF (PeproTech, AF-100-18B) and 20 ng/ml EGF (PeproTech, AF-100-15). For further passaging, cells were washed with PBS and detached with Accutase (Sigma-Aldrich, A6964) diluted 1:2 to 1:3 in PBS. Information for each model can be found in **Extended Data Table 5**. All RMS primary cultures were regularly tested to ensure no mycoplasma contamination with the LookOut® Mycoplasma PCR-Detektions-Kit (Sigma-Aldrich, MP0035-1KT).

### Culture of cell lines

The cell lines Rh4 (RRID: CVCL_5916), Rh41 (RRID: CVCL_2176; Peter Houghton, Greehey Children’s Cancer Research Institute, San Antonio, TX), KFR (RRID: CVCL_S637; Jindrich Cinatl, Abteilung für paediatrische Tumor und Virusforschung, Frankfurter Stiftung für krebskranke Kinder, Frankfurt) and RMS (RRID: CVCL_W527; Janet Shipley, Sarcoma Molecular Pathology, The Institute of Cancer Research) were cultured on uncoated plates in Dulbecco’s Modified Eagle’s Medium (DMEM medium) (Sigma Aldrich, D5671) supplemented with 10% fetal bovine serum (Life Technologies), 100 U/ml penicillin/streptomycin (Thermofisher, 15140122) and 2 mM Glutamax (Life technologies, 335050-061). For further passaging, cells were washed with PBS and detached with Trypsin (BioConcept, 5-51F00-I). All cell lines were authenticated by short tandem repeat analysis (STR) profiling and regularly tested to ensure no mycoplasma contamination with the LookOut® Mycoplasma PCR-Detektions-Kit (Sigma-Aldrich, MP0035-1KT)

### Doxycycline-inducible PAX3-FOXO1 knockdown

Rh4 and KFR cells containing doxycycline-inducible short hairpin RNAs (shRNAs) directed against P3F1 (shP3F1) or against a control hairpin (shSCR) were previously established as described^40^. ShRNA expression was induced using 50 ng/ml doxycycline in Rh4 cells and 10 ng/mL doxycycline in KFR cells.

### Sample preparation for single-cell RNA-sequencing

Dissociated PDX tumor cells were cultured as primary cultures for a low number of passages before sequencing (**Extended Data Table 5**). To study single-cell responses upon PAX3-FOXO1 downregulation, we induced shSCR or shP3F1 expression in Rh4 and KFR cells for 48 hrs as described above. For sequencing, cells were detached from the plates and washed 1x with PBS. Cells from different primary cultures or cell lines or experimental conditions were independently stained with a different oligonucleotide-tagged antibody (Totalseq^TM^-B hashtag antibodies) according to the manufacturer’s protocol and pooled together in equal proportion before sequencing in a single lane (**Extended Data Table 6**). In brief, 1-2 M cells were resuspended in 50 uL Cell Staining Buffer (Biolegend, 420201) and blocked with 5 µl of Human TruStain FcX™ (Biolegend, 422301) for 10 min at 4°C. Cells were then stained with 1.5 ug of Totalseq^TM^-B hashtag antibodies (Biolegend, 394631, 394633; 394635, 394637, 394639, 394641, 394643, 394645) in a total volume of 100 uL Cell Staining Buffer (Biolegend, 420201). After 30 min incubation on ice, cells were washed 3 times, pooled together, and resuspended in medium at a final concentration of 1000 cells/μl. Cells were stained with trypan blue and counted manually with a hemocytometer to determine their concentration. Viability was confirmed to be >85% before loading onto chip.

### Library preparation and sequencing

Cells were processed for library preparation according to the 10X Genomics Chromium Single Cell 3′ v3 workflow. Cell volume was adjusted to yield a recovery of ∼10,000 cells and loaded onto the 10X Genomics Single Cell A Chip. Library quality and concentration were assessed using High-Sensitivity D5000 ScreenTape (Agilent), and then sequenced on the Illumina NovaSeq 6000 system according to 10X Genomics recommendations.

### ScRNAseq hashtag sample de-multiplexing

Illumina sequencing reads were processed (de-multiplexing features and cellular barcodes, read alignment to the reference human genome GRCh38, and gene expression matrix generation) using the 10X Genomics Cell Ranger software Suite (version 3.0.1). Each gene expression matrix was then analyzed independently using Seurat R package (version 3.2.2)^28^ in R (version 3.6.1), with some modifications to the standard pipeline. We first de-multiplexed gene expression matrices based on hashtag oligonucleotide (HTOs) enrichment^45^. To do so, we added HTO data as a new independent assay to the RNA data and used centered log-ratio (CLR) transformation to normalize HTO raw counts. Cells were de-multiplexed using Seurat’s *HTODemux* function with default parameters. We assigned sample identities based on the maximal HTO signal and on the antibody sequence used for staining (**Supplementary Table 5**).

### ScRNAseq data analysis

De-multiplexed samples were analyzed independently. Low quality cells, identified as cells with <200 or >8000 genes, total number of transcripts (UMIs) <1000 or >50000, and/or percentage UMIs mapping to mitochondrial genes >15%, were removed. After log-normalizing the data, we assigned cell cycle scores per single-cell, using Seurat’s *CellCycleScoring* function, and scaled the expression of each gene regressing out the number of UMIs and the percentage UMIs mapping to mitochondrial genes. We then performed principal component analysis (PCA) to denoise the data and, based on elbow plots and the percentage of explained variance, selected the number of principal components (PCs) to consider for downstream analysis. We then built a shared nearest neighbor graph and used the Louvain algorithm for clustering (resolution of 0.2-0.3) the cell subpopulation. Single cells were visualized as uniform manifold approximation and projection (UMAP) plots. Differential expression analysis was performed using Seurat’s *FindAllMarkers* function only considering genes with >log_2_(0.25) fold-change and expressed in at least 5% of cells in the cluster. We annotated each clusters in the different datasets independently. Finally, to distinguish high-cycling from low-cycling cells, we used S and G2M phase scores calculated by the *CellCycleScoring* function. High-cycling cells were defined as cells with high S or G2M scores (>0), and low-cycling cells as the ones with low S and G2M scores (<0).

### ScRNAseq dataset integration

We performed four different dataset integrations [all RMS (*1*), aRMS primary cultures (*2*), eRMS primary cultures (*3*), aRMS cell lines (*4*)]. To integrate all RMS (*1*), we first reduced the data using reciprocal PCA (RPCA), given the large size of the dataset, and integrated them using the *FindIntegrationAnchors* function in Seurat with *k* = 5 neighbors when picking anchors. To integrate samples from aRMS (*2*) or eRMS primary cultures (*3*) or aRMS cell lines (*4*), we first transformed the data using *SCTransform* function, regressing out the number of UMI and the percentage of mitochondrial genes, and then integrated them using the *FindIntegrationAnchors* function in Seurat. Integrated data were then analyzed as described above.

### ScRNAseq pseudotime trajectory analysis

Pseudotime ordering of aRMS or eRMS was performed on a subset of cells labelled as *MuSC-like*, *cycling progenitors*, *S-phase* and *differentiated* from the integrated aRMS (*2*) or eRMS primary culture (*3*) datasets. We reduced and visualized the combined datasets using PHATE^39^ (*t* = 50, *knn* = 20) instead of UMAP, and used the *slingshot* package^38^ to organize cells in pseudotime and infer a trajectory. We set *MuSC-like* cells as the starting cluster for the trajectory calculation.

### ShP3F1 scRNAseq analysis

After hashtag de-multiplexing as described above, samples derived from Rh4 or KFR cells were merged and analyzed independently. We first log-normalizing the data, and then assigned myogenic program scores by using the gene signatures previously identified for *MuSC-like*, *cycling progenitors*, *S-phase* and *differentiated* clusters using Seurat’s *FindAllMarkers* function on the integrated aRMS cell line dataset (**Supplementary Table 1**). We then scaled the expression of each gene regressing out the number of UMI and the percentage of mitochondrial genes. We then performed principal component analysis (PCA) to denoise the data and, based on elbow plots, selected the number of principal components (PCs) to retain for downstream analysis. We then visualized single cells as UMAP plots. To identify genes differentially expressed upon P3F1 downregulation, we used Seurat’s *FindMarkers* function across shP3F1 cells cultured in the presence or absence of doxycycline (**Supplementary Table 2**).

### ScRNAseq mapping of aRMS cells to regenerating muscle cells

A subset of cells labelled as *MuSC-like*, *cycling progenitors*, *S-phase* and *differentiated* from the integrated aRMS primary culture dataset was mapped to a publicly available mouse single-cell dataset of regenerating muscle (GEO: GSE143437)^31^ using RPCA and normalized using *SCTransform* as described above. Mouse genes were first converted to human orthologs using the biomaRt R package^46^.

### RMS patient gene expression

To measure myogenic gene expression in RMS patients, we analyzed a published RMS patient gene expression dataset (Davicioni - 147 - MAS5.0 - u133a) available on the R2: Genomics Analysis and Visualization Platform (http://r2.amc.nl)^34^. We computed expression of PAX7, MYOD1, MYOG and MYH8 genes subsetting across aRMS or eRMS patients.

### Clinical outcomes of subpopulation signatures

To assess the clinical value of the subpopulations identified by scRNAseq, we first generated cluster-associated gene signatures by pairwise comparisons between all the clusters identified by scRNAseq in the combined aRMS primary culture dataset. Subpopulation-specific markers were identified using the likelihood-ratio test^47^ implemented in the *FindAllMarkers* function in Seurat and defined as genes overexpressed with a log-fold change > 0.25 and *P* < 0.05 following Bonferroni correction. Next, we analyzed the published RMS patient (Davicioni - 147 - MAS5.0 - u133a) gene expression dataset^34^ using the R2: Genomics Analysis and Visualization Platform (http://r2.amc.nl). For each gene signature, we computed the average log-fold change in expression between living (status = live) and deceased (status = dead) patients, where the averaging is over all the genes in the signature. Here, we consider both upregulated and downregulated genes. However, the downregulated genes contribute to the average log-fold change with minus signs. To obtain a null distribution for this test statistic, we randomly flip the “survivor” and “non-survivor” labels when computing average log-fold change. We simulate this null distribution to compute the *p*-value for each gene signature.

### Antibody conjugation with metal isotopes for mass cytometry

Purified antibodies were conjugated to the indicated metals for mass cytometry analysis using the MaxPAR X8 antibody labeling kit (Fluidigm) according to the manufacturer’s instruction. Following labeling, antibodies were diluted in Candor PBS Antibody Stabilization solution (Candor Bioscience GmbH, Wangen, Germany) to 0.2 mg/mL and stored long-term at 4°C. Each antibody clone and lot was titrated to optimal staining concentrations using human myoblasts.

### Mass cytometry sample preparation and staining

Cells from primary cultures (aRMS-1, aRMS-2, aRMS-6, aRMS-8, aRMS-9, eRMS-1.1, eRMS-1.2, eRMS-2.1, eRMS-4, eRMS-8.2, eRMS-8.3) or cell lines (Rh4, KFR, RMS) were first detached from plates and pulsed with IdU (Sigma Aldrich, I7125) at a final concentration of 50 μM for 30 min at 37°C. Dead cells were stained using cisplatin as previously described^48^. After washing with PBS, cells were resuspended in serum-free DMEM medium and cisplatin (Sigma Aldrich, P4394) was added for 1 min at a final concentration of 25 μM at room temperature. Reaction was quenched by adding 3 mL DMEM medium with 10% FBS. Cells were then centrifuged at 1200 rpm for 10 min at 4 °C, resuspended in cell staining media (CSM: PBS with 0.5% bovine serum albumin (BSA: Sigma Aldrich, A8022) and 0.02% sodium azide) and fixed with 1.6% paraformaldehyde (PFA: ThermoFisher, 28908) for 10 minutes on ice. Cells were washed with CSM and stained with antibodies against surface markers included in the mass cytometry panel for 1h at room temperature. Cells were washed twice with CSM and permeabilized with methanol for 10 minutes on ice. Cells were washed twice with CSM and stained with antibodies against intracellular markers included in the mass cytometry panel for 1h at room temperature. Cells were washed twice with CSM and stained with 1 mL of 191/193Ir DNA intercalator (Fluidigm) diluted in PBS (1:5000) with 1.6% PFA for 20 mins at room temperature.

### Mass cytometry measurement

Cells were acquired on the CyTOF2 mass cytometer (Fluidigm) at an event rate of approximately 500 cells per second as previously described^35^. The instrument was run in high-resolution mode (Mass resolution ∼700) with internally calibrated dual-count detection. Noise reduction and cell extraction parameters were: cell length 10-65, lower convolution threshold 10. Samples were normalized using beta beads^49^.

### X-Shift analysis and graphic display of single-cell mass cytometry data

Pooled aRMS and pooled eRMS cells were clustered based on a combination of surface markers, myogenic transcription factors and cell-cycle markers (CD44, Axl, Pax-7, myogenin, IdU) using the X-Shift algorithm. In order to visualize the spatial relationships between cells within these X-shift clusters, 2000 randomly sampled cells from each cluster were subjected to a force-directed layout^36^. All conditions were processed simultaneously so that the resulting map would capture all populations present in the entire dataset.

### Immunofluorescence

To assess CD44, Pax-7, myogenin or MyHC positivity, cells were first washed with PBS and then fixed with ROTI®Histofix 4 % (Carl Roth, P087.3) for 30 min. Following 3x5 min washes with PBS using an automated plate washer (BioTek 50 TS washer, Agilent Technologies), cells were permeabilized with 0.1% Triton-X 100 (Sigma Aldrich, X100-100ML) in PBS for 15 min. After 3x5 min washes with PBS and blocking with 1% bovine serum albumin (BSA: Sigma Aldrich, A8022) in PBS for 1 hour, cells were incubated for 2 hrs at room temperature or overnight at 4°C with primary antibody diluted in 1% BSA. Cells were washed 3x5 min with PBS and incubated with secondary antibodies diluted 1:300 in 1% BSA. Finally, cells were washed again 3x5 min with PBS and covered with 1:1000 solution of Hoechst 33342 (ThermoFisher Scientific, 62249) in PBS. Images were acquired and quantified with an automated workflow on PerkinElmer Operetta. Primary antibodies used include: myogenin (M-225: Santa Cruz, sc-576 or ThermoFisher, PA5-78067; dilution: 1:300), MyHC (MF-20: Developmental Studies Hybridoma Bank; dilution: 1:500), Ki67 (ThermoFisher, 14-5698-82; dilution: 1:1000), FITC anti-human CD44 Antibody (Biolegend, 338804; dilution: 1:100). Secondary antibodies used included: Goat anti-Mouse IgG (H+L) Highly Cross-Adsorbed Secondary Antibody Alexa Fluor 594 (ThermoFisher, A-11032), Goat anti-Rabbit IgG (H+L) Highly Cross-Adsorbed Secondary Antibody Alexa Fluor 488 (ThermoFisher, A-11034), Goat anti-Rat IgG (H+L) Highly Cross-Adsorbed Secondary Antibody Alexa Fluor 647 (ThermoFisher, A21247) at a dilution of 1:1000.

### MYOscopy

To assess the cellular composition of aRMS cells based on markers identified by single-cell analyses, cells were plated in 384-well plates and processed for IF as described above with the following primary antibodies: myogenin (M-225: Santa Cruz, sc-576 or ThermoFisher, PA5-78067; dilution: 1:300), MyHC (MF-20: Developmental Studies Hybridoma Bank; dilution: 1:500) and Ki-67 (ThermoFisher, 14-5698-82; dilution: 1:1000). Secondary antibodies used included: Goat anti-Mouse IgG (H+L) Highly Cross-Adsorbed Secondary Antibody Alexa Fluor 594 (ThermoFisher, A-11032), Goat anti-Rabbit IgG (H+L) Highly Cross-Adsorbed Secondary Antibody Alexa Fluor 488 (ThermoFisher, A-11034), Goat anti-Rat IgG (H+L) Highly Cross-Adsorbed Secondary Antibody Alexa Fluor 647 (ThermoFisher, A21247) at a dilution of 1:1000.

### Western blotting

Whole cell lysates were extracted using RIPA lysis buffer (50 mM Tris-HCl pH 7.5, 150 mM NaCl, 1% NP-40, 0.5% Na-deoxycholate, 0.1% SDS, 1 mM EGTA, 50 mM NaF, 5 mM Na_4_P_2_O_7_, 1 mM Na_3_VO_4_, and 10 mM ß-glycerol phosphate) in the presence of the cOmplete^™^, Mini, EDTA-free Protease Inhibitor Cocktail (Sigma Aldrich, 11836170001). Protein concentration was measured with Pierce^TM^ BCA Protein Assay Kit (Thermo Fisher Scientific, 23227) according to the manufacturer’s instructions. 5-20 ug of whole cell lysates were reduced with Laemmli Sample Buffer (Bio-rad, 161-0747) supplemented with 1:20 DTT. After boiling the samples at 95°C for 5 minutes, proteins were separated using NuPAGE™ 4 to 12%, Bis-Tris, 1.0 mm, Mini Protein Gels (Thermo Fisher Scientific, NP0323BOX). Gels were transferred on membranes using Trans-Blot® Turbo™ Transfer System (Bio-rad, 1704150). Following blocking with 5% milk (Carl Roth, T145.3) in TBS/0.05% tween (Sigma Aldrich, P9416) for 30-60 min, membranes were incubated overnight at 4°C or for 2 hrs at room temperature (RT) with primary antibodies diluted 1:1000. After three washing steps with TBS/0.05% tween for 5 min, membranes were incubated for 1 hr at RT with HRP-linked secondary antibodies diluted 1:5000. Finally, after three additional washing steps with TBS/0.05% tween for 5 min, proteins were detected by chemiluminescence using SuperSignal™ West Femto Maximum Sensitivity Substrate (Thermo Fisher Scientific, 34095) and a ChemiDoc MP imager (BioRad). The following antibodies were used: Pax-7 (Developmental Studies Hybridoma Bank, DSHB), MyoD (M-318: Santa Cruz, sc-760), myogenin (M-225: Santa Cruz, sc-576 or ThermoFisher, PA5-78067), PAX3-FOXO1 (FKHR H-128: Santa Cruz, sc-11350), MyHC (MF-20: Developmental Studies Hybridoma Bank), GAPDH (Cell Signaling, 5174S), pERK (Cell Signaling, 9101S), ERK1/2 (Cell Signaling, 9102L), Anti-rabbit IgG HRP-linked Antibody (Cell Signaling, 7074S), Anti-mouse IgG HRP-linked Antibody (Cell Signaling, 7076S).

### Immunohistochemistry

Formalin-fixed, paraffin-embedded (FFPE) PDX tumors were first cut in sections of 2 μM and then stained on the BOND Fully Automated IHC Staining System (Leica). The sections were incubated for 30 min with primary antibodies against PAX7 (DSHB; dilution: 1:100), MYOG (Cell Marque Lifescreen, 296M-14; dilution: 1:20), MyHC (DSHB; dilution: 1:500), Ki-67 (Cell Marque Lifescreen, 275R-16; dilution: 1:100). Visualization of the antibodies was performed with a Bond refine detection system, Leica. All sections were counterstained with haematoxylin.

### Flow cytometry and FACS sorting

CD44^+^ and CD44^-^ subpopulations were sorted on a BD FACS Aria Fusion using FITC anti-human CD44 antibody (Biolegend, 338804) at 1:200 dilution. CD44-positive and -negative cells were gated based on the FITC Mouse IgG1 (Biolegend, 400110) isotype control. To exclude dead cells from sorting, cells were stained with eBioscience™ 7-AAD Viability Staining Solution (ThermoFisher, 00-6993-50) prior to sorting. For stability experiments, 50’000 cells/well from CD44^+^, CD44^-^ or unsorted populations were cultured in 24-well plates and analyzed on a LSRFortessa (BD) at regular intervals for 3 weeks after fresh staining with FITC anti-human CD44 Antibody as described above.

To measure AXL, CD44 and CD105 expression after drug treatment, cells were treated with drugs for 48 hrs as described below and detached from the plates for staining. Cells were first stained with Zombie NIR™ Fixable Viability Kit (Biolegend, 423105) according to the manufacturer’s instructions to gate out dead cells. Flow cytometry was performed by labeling cells for 30 min at 4°C with the following antibodies (dilution of 1:50): FITC anti-human CD44 (Biolegend, 338804), PE anti-CD105 (Biolegend, 800503) and APC Axl (Thermo Fisher Scientific, 17-1087-41). Isotype controls were used to gate positive/negative cells and included the following antibodies: FITC Mouse IgG1 (Biolegend, 400110), APC Mouse IgG1 (Biolegend, 400122) and PE Mouse IgG1 (Biolegend, 400114). All FACS analysis were done on FlowJo™ v10.8 software.

### High-content single-cell imaging drug screening

aRMS-1 and aRMS-3 cells were plated in 384-well plates coated with Matrigel at a cell density of 2000-4000 cells/well. The day after, cells were treated with the drug library containing 244 drugs at a concentration of 10 μM, 1 μM, 0.1 μM or 0.01 μM. After 72 hrs incubation, cells were processed for MYOscopy as described above. Cells were assigned to the corresponding myogenic states according to expression of the following markers: *quiescent MuSC-like* (myogenin^-^Ki-67^-^), *cycling MuSC-like* (myogenin^-^Ki-67^+^), *cycling progenitors* (myogenin^+^Ki-67^+^MyHC^-^), *non-cycling committed progenitors* (myogenin^+^Ki-67^-^ MyHC^-^), *differentiated* (MyHC^+^). An unstained control was included on every plate and used as a reference for signal background. Differentiating hits were defined as drugs increasing the percentage of *non-cycling committed progenitors* and *differentiated* cells and decreasing *cycling progenitors* compared to untreated controls; de-differentiating hits were defined as drugs increasing the percentage of *MuSC-like* cells compared to untreated controls of at least 1.5-fold. An overall “differentiating score” was calculated as follows: *[% non-cycling committed progenitors (myogenin^+^Ki-67^-^MyHC^-^) + % differentiated (MyHC^+^) - % cycling progenitors (myogenin^+^Ki-67^+^MyHC^-^))]_compound, 1 µM_ - [% committed (myogenin^+^Ki-67^-^MyHC^-^) + % differentiated (MyHC^+^) - % cycling progenitors (myogenin^+^Ki-67^+^MyHC^-^)_untreated control_*; whereas an overall “de-differentiating score” was calculated as follows: *[% quiescent MuSC-like (myogenin^-^Ki67^-^) + % cycling MuSC-like (myogenin ^-^Ki67^+^)]_compound, 1 µM_ - [% quiescent MuSC-like (myogenin ^-^Ki67^-^) + % cycling MuSC-like (myogenin ^-^Ki67^+^)]_untreated control_*.

For the trametinib-combination screening, aRMS-1 cells were plated in 384-well plates coated with Matrigel at a cell density of 4000 cells/well. The day after, cells were treated with the drug library containing 244 drugs at a concentration of 10 μM, 1 μM, 0.1 μM or 0.01 μM and with additional 50 nM trametinib (Selleckchem, S2673). As a control, untreated cells were included on every plate. After 72 hrs incubation, cells were processed for immunofluorescence as described above. The “combined differentiating score” was calculated averaging the “differentiating scores” calculated at 10 µM, 1 µM, 100 nM and 10 nM and subtracting the differentiating score of 50 nM trametinib alone.

### Drug treatment

For WB, qRT-PCR and cell cycle analysis, cells were plated in 6-well plates at a concentration of 300’000 cells/well, equilibrated overnight and then treated with drugs for the indicated time. In case of trametinib, cells were treated immediately the day of plating. For WB analysis of phosphorylated ERK, cells were plated in 6-well plates at a concentration of 300’000 cells/well, equilibrated overnight and then treated with 5 or 10 nM Trametinib for 3 hrs.

For flow cytometry analysis, cells were plated in 24-well plates at a concentration of 100’000 cells/well, equilibrated overnight and then treated with chemotherapy for 48 hrs.

For immunofluorescence, cells were plated in 384-well plates at a concentration of 2000- 4000 cells/well, equilibrated overnight, and then treated with the indicated drugs for 72 hrs. To test the drug sensitivity of CD44^+^ and CD44^-^ FACS-sorted subpopulations, cells were sorted as described above. After sorting, CD44^+^ and CD44^-^ subpopulations, together with an unsorted reference population, were plated in 384-well plates coated with Matrigel at a cell density of 800 cells/well. The day after, the medium was replaced, and cells were incubated for further 72 hrs with the indicated drugs. Data of each CD44^+^, CD44^-^ and unsorted populations were normalized to DMSO (vehicle)-treated conditions, defined as 100% viability. IC_50_ values were determined from the dose-response curves generated using GraphPad Prism.

Drugs included vincristine sulfate (ApexBio, A1765-5.1), 4-Hydroperoxycyclophosphamide (4-HC: Niomech, CAS 39800-16-3), etoposide (Selleckchem, S1225) and trametinib (Selleckchem, S2673).

### *In vivo* PDX treatment

Dissociated aRMS-1 cells were first expanded *in vitro* for three passages. Two million aRMS-1 cells were injected orthotopically into the gastrocnemius muscle of NSG mice. Treatment was started when tumors became palpable (day 17 after injection) for two weeks. For trametinib treatment (*n* = 3 mice), drug (1 mg/kg; Selleckchem, S2673) was dissolved in PBS+1% DMSO and administered by intraperitoneal injection (i.p.) five times a week. For dabrafenib treatment (*n* = 3 mice), drug (15 mg/kg; Selleckchem, S2807) was dissolved in PBS+5% DMSO and administered by oral gavage 5 times a week. The combination group (*n* = 3 mice) was treated as described above with both trametinib and dabrafenib. Control mice were treated by i.p. (*n* = 3) or by oral gavage (*n* = 2) with the vehicles PBS+1% DMSO, respectively PBS+5% DMSO. Mice were euthanized at the end of the second treatment week (day 28 after injection). Tumors were harvested four hours after the last treatment and processed for qRT-PCR and IHC.

For vincristine sulfate treatment (*n* = 3 mice), drug (10 mg/kg; MedChemExpress, HY-N0488) was dissolved in PBS and administered by i.p. injection two times a week (Monday and Thursday). A mouse treated with vincristine was euthanized at day 18 after injection due to severe body weight loss (>20% than baseline) and was therefore excluded from the analysis; the other *n* = 2 mice were euthanized at the end of the first, respectively of the second, week after the start of the treatment (day 21 and day 28).

### Real-Time Quantitative Reverse Transcription PCR (qRT-PCR)

Total RNA was extracted using the RNeasy Micro Kit (Qiagen, 74004) from cultured cells and using the RNeasy Mini Kit (Qiagen, 74106) from tumor pieces according to manufacturer’s instructions. cDNA was synthesized using the High-Capacity cDNA Reverse Transcription Kit with RNase Inhibitor (Thermo Fisher Scientific, 4374967) modifying the total volume to 10 μL. In specific, for each sample, a mix consisting of 1.0 μL 10X RT Buffer, 0.4 μL 25XdNTPs Mix (100 mM), 1.0 μL 10X RT Random Primers, 0.5 μL MultiScribeTM Reverse Transcriptase, 0.5 μL RNase inhibitor, 1.6 μL Nuclease-free H_2_O and 5 μL Nuclease-Free Water (Thermo Fisher Scientific, AM9937) was incubated in a thermocycler according to manufacturer’s instructions. Each sample was analyzed by qRT-PCR using TaqMan™ Gene Expression Master Mix (Thermofisher Scientific, 4369542) and TaqMan gene expression assays. QRT-PCR was performed in technical triplicates for each sample on a 7900HT Fast Real-Time PCR machine. To calculate the relative gene expression of each gene, the 2^-ΔΔCt^ method was used and the quantity of RNA was normalized to the internal control GAPDH. The following TaqMan gene expression assays were used: CD44 (Hs01075864_m1), AXL (Hs01064444_m1), MYOD1 (Hs00159528_m1), MYOG (Hs01072232_m1), MYH3 (Hs01074230_m1), GAPDH (Hs02758991_g1).

### Cell cycle analysis

After treatment with Trametinib as described above, cells were detached from the plates, washed in PBS, fixed in ice-cold 70% ethanol and incubated at −20 °C for at least 2 h. Cells were then washed with PBS and resuspended in 500 μl PI solution (0.4 mg/ml PI and 0.4 μg/ml RNase A in PBS with 0.0001% Triton X-100) before analysis on a LSRFortessa (BD) Cell Analyzer. All FACS analysis were done on FlowJo™ v10.8 software.

### Data availability

All sequencing datasets will be made available upon manuscript acceptance or request on the Gene Expression Omnibus (GEO) database. Codes will be shared upon reasonable request to the corresponding author.

## ACKNOWLEDGEMENTS

We thank: staff from the Functional Genomics Center Zürich (FGCZ), in particular Dr. Emilio Yángüez, Dr. Doris Popovic, Andreia Cabral de Gouvea, for support with the single-cell RNA sequencing; Susanne Dettwiler and Fabiola Prutek from the Institut für Pathologie und Molekularpathologie (Universitätsspital Zürich) for processing and staining the histology sections from the PDXs; Dr. Quy Ngo for initial support with the single-cell RNA sequencing data analysis.

Research reported in this publication was supported by: Cancer League Switzerland (KLS-5143-08-2020) to B.W.S.; the Childhood Research Foundation Switzerland to B.W.S.; the Swiss National Science Foundation (3100-175558) to B.W.S.; the Children’s Research Center grant from the University Children’s Hospital Zurich (project number: 10788_E) to S.G.D; the California Institute for Regenerative Medicine grant (RB5-07469) to H.M.B.; the Baxter Foundation to H.M.B. and the BD Biosciences Stem Cell grant to E.P.

## REFERENCES

1. Tirosh, I. & Suvà, M. L. Deciphering human tumor biology by single-cell expression profiling. Annual Review of Cancer Biology (2019).

2. Hanahan, D. Hallmarks of Cancer: New Dimensions. Cancer Discov. 12, 31–46 (2022).

3. Filbin, M. & Monje, M. Developmental origins and emerging therapeutic opportunities for childhood cancer. Nat. Med. 25, 367–376 (2019).

4. Filbin, M. G. et al. Developmental and oncogenic programs in H3K27M gliomas dissected by single-cell RNA-seq. Science (80-.). (2018).

5. Zhang, L. et al. Single-Cell Transcriptomics in Medulloblastoma Reveals Tumor-Initiating Progenitors and Oncogenic Cascades during Tumorigenesis and Relapse. Cancer Cell (2019).

6. Couturier, C. P. et al. Single-cell RNA-seq reveals that glioblastoma recapitulates a normal neurodevelopmental hierarchy. Nat. Commun. (2020).

7. Gojo, J. et al. Single-Cell RNA-Seq Reveals Cellular Hierarchies and Impaired Developmental Trajectories in Pediatric Ependymoma. Cancer Cell (2020).

8. Skapek, S. X. et al. Rhabdomyosarcoma. Nature Reviews Disease Primers (2019).

9. Hettmer, S. & Wagers, A. J. Muscling in: Uncovering the origins of rhabdomyosarcoma. Nature Medicine (2010).

10. Charytonowicz, E., Cordon-Cardo, C., Matushansky, I. & Ziman, M. Alveolar rhabdomyosarcoma: Is the cell of origin a mesenchymal stem cell? Cancer Lett. 279, 126–136 (2009).

11. Langenau, D. M. et al. Effects of RAS on the genesis of embryonal rhabdomyosarcoma. Genes Dev. 21, 1382–1395 (2007).

12. Keller, C. & Capecchi, M. R. New genetic tactics to model alveolar rhabdomyosarcoma in the mouse. Cancer Res. 65, 7530–7532 (2005).

13. Drummond, C. J. et al. Hedgehog Pathway Drives Fusion-Negative Rhabdomyosarcoma Initiated From Non-myogenic Endothelial Progenitors. Cancer Cell 33, 108–124.e5 (2018).

14. Hatley, M. E. et al. A Mouse Model of Rhabdomyosarcoma Originating from the Adipocyte Lineage. Cancer Cell 22, 536–546 (2012).

15. Pappo, A. S. et al. Survival After Relapse in Children and Adolescents With Rhabdomyosarcoma: A Report From the Intergroup Rhabdomyosarcoma Study Group. J. Clin. Oncol. 17, 3487–3493 (1999).

16. Heske, C. M. & Mascarenhas, L. Relapsed Rhabdomyosarcoma. J. Clin. Med. 10, 804 (2021).

17. Shern, J. F. et al. Genomic Classification and Clinical Outcome in Rhabdomyosarcoma: A Report From an International Consortium. J. Clin. Oncol. 39, 2859–2871 (2021).

18. Shern, J. F. et al. Comprehensive genomic analysis of rhabdomyosarcoma reveals a landscape of alterations affecting a common genetic axis in fusion-positive and fusion-negative tumors. Cancer Discov. (2014).

19. Wachtel, M. & Schäfer, B. W. PAX3-FOXO1: Zooming in on an “undruggable” target. Semin. Cancer Biol. 50, 115–123 (2018).

20. Regina, C. et al. Negative correlation of single-cell PAX3:FOXO1 expression with tumorigenicity in rhabdomyosarcoma. Life Sci. Alliance 4, 1–18 (2021).

21. Kikuchi, K. et al. Cell-Cycle Dependent Expression of a Translocation-Mediated Fusion Oncogene Mediates Checkpoint Adaptation in Rhabdomyosarcoma. PLoS Genet. 10, (2014).

22. Boudjadi, S. et al. A fusion transcription factor⇓driven cancer progresses to a fusion-independent relapse via constitutive activation of a downstream transcriptional target. Cancer Res. 81, 2930–2942 (2021).

23. Cruz, F. Dela & Matushansky, I. Solid tumor differentiation therapy - Is it possible? Oncotarget 3, 559–567 (2012).

24. De Thé, H. Differentiation therapy revisited. Nat. Rev. Cancer 18, 117–127 (2018).

25. Garcia, N. et al. Vertical Inhibition of the RAF-MEK-ERK Cascade Induces Myogenic Differentiation, Apoptosis and Tumor Regression in H/NRAS Q61X-mutant Rhabdomyosarcoma. Mol. Cancer Ther. (2021).

26. Yohe, M. E. et al. MEK inhibition induces MYOG and remodels super-enhancers in RAS-driven rhabdomyosarcoma. Sci. Transl. Med. 10, 1–18 (2018).

27. Fox, E. J. & Loeb, L. A. One cell at a time. Nat. News Views 512, 143–144 (2014).

28. Stuart, T. et al. Comprehensive Integration of Single-Cell Data. Cell 177, 1888–1902.e21 (2019).

29. Zhou, Y. et al. Metascape provides a biologist-oriented resource for the analysis of systems-level datasets. Nat. Commun. 10, (2019).

30. Oprescu, S. N., Yue, F., Qiu, J., Brito, L. F. & Kuang, S. Temporal Dynamics and Heterogeneity of Cell Populations during Skeletal Muscle Regeneration. iScience 23, 100993 (2020).

31. De Micheli, A. J. et al. Single-Cell Analysis of the Muscle Stem Cell Hierarchy Identifies Heterotypic Communication Signals Involved in Skeletal Muscle Regeneration. Cell Rep. (2020).

32. Tirosh, I. et al. Dissecting the multicellular ecosystem of metastatic melanoma by single-cell RNA-seq. Science (80-.). 352, 189–196 (2016).

33. R2: Genomics Analysis and Visualization Platform. Available at: http://r2.amc.nl.

34. Davicioni, E. et al. Identification of a PAX-FKHR gene expression signature that defines molecular classes and determines the prognosis of alveolar rhabdomyosarcomas. Cancer Res. 66, 6936–6946 (2006).

35. Porpiglia, E. et al. High-resolution myogenic lineage mapping by single-cell mass cytometry. Nat. Cell Biol. 19, 558–567 (2017).

36. Samusik, N., Good, Z., Spitzer, M. H., Davis, K. L. & Nolan, G. P. Automated mapping of phenotype space with single-cell data. Nat. Methods 13, 493–496 (2016).

37. Behbehani, G. K., Bendall, S. C., Clutter, M. R., Fantl, W. J. & Nolan, G. P. Single-cell mass cytometry adapted to measurements of the cell cycle. Cytom. Part A 81 A, 552–566 (2012).

38. Street, K. et al. Slingshot: cell lineage and pseudotime inference for single-cell transcriptomics. BMCGenomics 19, 1–16 (2018).

39. Moon, K. R. et al. Visualizing structure and transitions in high-dimensional biological data. Nat. Biotechnol. 37, 1482–1492 (2019).

40. Ommer, J. et al. Aurora a kinase inhibition destabilizes PAX3-FOXO1 and MYCN and synergizes with navitoclax to induce rhabdomyosarcoma cell death. Cancer Res. 80, 832–842 (2020).

41. Knispel, S. et al. The safety and efficacy of dabrafenib and trametinib for the treatment of melanoma. Expert Opin. Drug Saf. 17, 73–87 (2018).

42. Kelly, R. J. Dabrafenib and trametinib for the treatment of non-small cell lung cancer. Expert Rev. Anticancer Ther. 18, 1063–1068 (2018).

43. Ebauer, M., Wachtel, M., Niggli, F. K. & Schäfer, B. W. Comparative expression profiling identifies an in vivo target gene signature with TFAP2B as a mediator of the survival function of PAX3/FKHR. Oncogene 26, 7267–7281 (2007).

44. Manzella, G. et al. Phenotypic profiling with a living biobank of primary rhabdomyosarcoma unravels disease heterogeneity and AKT sensitivity. Nat. Commun. (2020).

45. Stoeckius, M. et al. Cell Hashing with barcoded antibodies enables multiplexing and doublet detection for single cell genomics. Genome Biol. 19, 1–12 (2018).

46. Durinck, S., Spellman, P. T., Birney, E. & Huber, W. Mapping identifiers for the integration of genomic datasets with the R/ Bioconductor package biomaRt. Nat. Protoc. 4, 1184–1191 (2009).

47. McDavid, A. et al. Data exploration, quality control and testing in single-cell qPCR-based gene expression experiments. Bioinformatics 29, 461–467 (2013).

48. Fienberg, H. G., Simonds, E. F., Fantl, W. J., Nolan, G. P. & Bodenmiller, B. A platinum-based covalent viability reagent for single-cell mass cytometry. Cytom. Part A 81 A, 467–475 (2012).

49. Finck, R. et al. Normalization of mass cytometry data with bead standards. Cytom. Part A 83 A, 483–494 (2013).

