## Extended Data Table for "Single-cell mapping of tumor heterogeneity in pediatric rhabdomyosarcoma reveals developmental signatures with therapeutic relevance"

Extended Data Tables

### **Extended Data Table 1:** PDX characteristics and information on the clinical status of the patients.

| PDX | Alternative name | Sex | Age | RMS subtype | Site | Status | Location | Tumor treatment | Hospital of origin |
| --- | --- | --- | --- | --- | --- | --- | --- | --- | --- |
| aRMS-1 | IC-pPDX-104 | F | 7 | aRMS | Primary | Recurrent | Tibia | Yes | Paris |
| aRMS-2 | IC-pPDX-29 | F | 14 | aRMS | Primary | Recurrent | Paravertebral | Yes | Paris |
| aRMS-3 | IC-pPDX-35 | M | 13 | aRMS | Metastatic | Recurrent | mediastinum | Yes | Paris |
| aRMS-4 | SJRHB013759_X1 (Mast118) | M | 18 | aRMS | Metastatic | Recurrent | Inguinal | No | St. Jude |
| aRMS-5 | Berlin 13304 |  |  | aRMS | NA | NA | NA | NA | Berlin |
| aRMS-6 | RMS-ZH-003 | F | 3 | aRMS | NA | Recurrent | Paraurethral | NA | Zürich |
| aRMS-7 | Berlin 11410 | F | 10 | aRMS | Primary | Diagnostic | Hand | No | Berlin |
| aRMS-8 | Berlin 14419 | F | 16 | aRMS | Primary | Recurrent | Forefoot | Yes | Berlin |
| aRMS-9 | Berlin 10752 | M | 12 | aRMS | Primary | Diagnostic | Forefoot | No | Berlin |
| aRMS-10 | RMS-Wi-001B | M | 23 | aRMS | Metastatic | Recurrent | Inguinal | Yes | Zürich |
| aRMS-11 | SJRHB010463_X16 (Mast60) | M | 18 | aRMS | Metastatic | Recurrent | Lung | No | St. Jude |
| eRMS-1.1 | SJRHB011_Y (Rh70) | M | 4 | eRMS | Primary | Diagnostic | Skull | Yes | St. Jude |
| eRMS-1.2 | SJRHB011_X (Rh73) | M | 5 | eRMS | Primary | Recurrent | Neck | No | St. Jude |
| eRMS-2.1 | SJRHB012_X (Rh71) | M | 17 | eRMS | Primary | Diagnostic | Prostate | No | St. Jude |
| eRMS-2.2 | SJRHB012_Y (Rh74) | M | 18 | eRMS | Primary | Recurrent | Prostate | Yes | St. Jude |
| eRMS-2.3 | SJRHB012_Z (Rh75) | M | 18 | eRMS | Primary | Recurrent | Prostate/bladder | Yes | St. Jude |
| eRMS-3.1 | SJRHB13758_X1  (Mast111) | F | 4 | eRMS | Primary | Diagnostic | Abdomen | No | St. Jude |
| eRMS-3.2 | SJRHB13758_X2  (Mast139) | F | 5 | eRMS | Primary | Recurrent | Abdomen | Yes | St. Jude |
| eRMS-4 | IC-pPDX-82 | F | 14 | eRMS | Primary | Diagnostic | Oral cavity | NA | Paris |
| eRMS-5 | Berlin 11410 | F | 10 | aRMS | Primary | Diagnostic | Hand | No | Berlin |
| eRMS-6 | Berlin 12181 | F | 5 | eRMS | Primary | Recurrent | Pararectal | Yes | Berlin |
| eRMS-7 | Berlin 14419 | F | 16 | aRMS | Primary | Recurrent | Forefoot | Yes | Berlin |
| eRMS-8.1 | Berlin 13454 | F | <1 | eRMS | Primary | Diagnostic | Head | No | Berlin |
| eRMS-8.2 | Berlin 13870 | F | <1 | eRMS | Primary | Recurrent | Head | Yes | Berlin |
| eRMS-8.3 | Berlin 13933 | F | 1 | eRMS | Primary | Recurrent | Head | Yes | Berlin |

M, male; F, female; NA, not available

**Extended Data Table 2:** Myogenic markers defined by De Micheli, A. J. *et al.* Single-Cell Analysis of the Muscle Stem Cell Hierarchy Identifies Heterotypic Communication Signals Involved in Skeletal Muscle Regeneration. *Cell Rep.* (2020). Genes marked with an “*****” indicate common expression in the RMS subpopulations.

| Quiescent MuSCs (Qu) | Cycling progenitors (Cy) | Committed progenitors (Co) |
| --- | --- | --- |
| CHODL | **STMN1*^2^** | SPG21 |
| CEBPB | **HMGB2*^2^** | IFFO1 |
| BTG2 | H2AFZ | **JAM3*^3^** |
| CRLF1 | **BIRC5*^2^** | **NEB*^3^** |
| GPX3 | **CENPA*^2^** | **CDKN1C*^3^** |
| JUND | **TUBA1B*^2^** | **ARPP21*^3^** |
| FOSB | **CKS2*^2^** | SVBP |
| EGR1 | RRM2 | S100A16 |
| NPPC | RAN | **ACTA2*^3^** |
| MAFF | DUT | OLFML2B |
| DNAJB1 | CKS1B | **ACTC1*^3^** |
| NR4A1 | HMGN2 | **TNNT2*^3^** |
| JUNB | HMGB1 | **TPM2*^3^** |
| ZFP36 | UBE2C | TMEM8C |
| **PPP1R15A*^1^** | TUBB5 | **MYL4*^3^** |
| HSPA1A | **CCNB2*^2^** | SLC29A1 |
| KLF4 | **CDK1*** | **TTN*^3^** |
| FOS | **TOP2A*^2^** | GM7325 |
| **CXCL1*^1^** | TK1 | **TNNI1*^3^** |
| **ERRFI1*^1^** | **CDC20*^2^** | **CKB*^3^** |
| MEG3 | **CENPF*^2^** | **CHRNA1*^3^** |
| MT1 | **SMC4*^2^** | **MYOG*^3^** |
| ID3 | 2700094K13RIK | C1QTNF3 |
| JUN |  | TMSB4X |
| SDC4 |  | SPP1 |
| PAX7 |  |  |

^1^Marker genes of RMS MuSC-like

^2^Marker genes of RMS cycling progenitors

^3^Marker genes of RMS differentiated cells

**Extended Data Table 3:** Myogenic markers defined by Oprescu, S. N., Yue, F., Qiu, J., Brito, L. F. & Kuang, S. Temporal Dynamics and Heterogeneity of Cell Populations during Skeletal Muscle Regeneration. *iScience* **23**, 100993 (2020). Genes marked with an “*****” indicate common expression in the RMS subpopulations.

| Quiescent MuSCs (Qsc) | Activated MuSCs (Asc) | Immunomyoblasts (Imb) | Dividing MuSCs (Div) | Committed MuSCs (Com) | Differentiated (Dif) |
| --- | --- | --- | --- | --- | --- |
| **Jund*^1^** | Fcerg1 | **Mest*^1^** | **Top2a*^2^** | Tmem8c | **Myh3*^3^** |
| Btg2 | C1qc | **Islr*^1^** | **Mki67*^2^** | **Myog*^3^** | **Tnni1*^3^** |
| Nppc | Ctss | Capn6 | **Cenpf*^2^** | Gm7352 | **Myl4*^3^** |
| Hspa1b | Tyrobp | Col1a2 | Hist1h1b | Ttn***^3^** | Acta1 |
| Atf3 | C1qa | **Plagl1*^1^** | 2810417H13Rik | Tnnt2***^3^** | **Myl1*^3^** |
| Fosb | C1qb | Crip1 | **Prc1*^2^** | Neb***^3^** | **Tnnt3*^3^** |
| Hspa1a | Lyz2 | Itm2a | **Stmn1*^2^** | Actc1***^3^** | **Mylpf*^3^** |
| Fos | **Ctsb*^1^** | **Mgp*^1^** | **Smc4*^2^** | Spg21 |  |
| Meg3 | Cd74 | **Igfbp5*^1^** | **Hmgb2*^2^** | Cdkn1c***^3^** |  |
| Jun | Apoe | Spp1 | **Ube2c*^2^** | Acta2***^3^** |  |

^1^Marker genes of RMS MuSC-like

^2^Marker genes of RMS cycling progenitors

^3^Marker genes of RMS differentiated cells

**Extended Data Table 4:** Number of cells in the identified Louvain clusters based on the status, location, tumor model, sample of origin, RMS subtype, treatment and cycling properties.

|  |  | S-phase | MuSc-like | Cycling progenitors | G1-phase | Differentiated |
| --- | --- | --- | --- | --- | --- | --- |
| **Status** | diagnostic | 1593 | 1007 | 688 | 631 | 771 |
|  | recurrent | 16628 | 7937 | 8162 | 7309 | 4133 |
| **Location** | NA | 688 | 175 | 385 | 298 | 319 |
|  | metastasis | 6430 | 1408 | 3019 | 2341 | 1421 |
|  | primary | 11103 | 7361 | 5446 | 5301 | 3164 |
| **Model** | Cell line | 5085 | 549 | 2368 | 1660 | 764 |
|  | PDC | 13136 | 8395 | 6482 | 6280 | 4140 |
| **Sample** | KFR | 1676 | 132 | 857 | 490 | 341 |
|  | Rh4 | 1851 | 163 | 650 | 652 | 195 |
|  | RMS | 1558 | 254 | 861 | 518 | 228 |
|  | aRMS-3 | 886 | 663 | 347 | 537 | 484 |
|  | aRMS-1 | 1064 | 275 | 588 | 372 | 354 |
|  | aRMS-2 | 924 | 335 | 334 | 496 | 585 |
|  | aRMS-4 | 459 | 196 | 304 | 144 | 173 |
|  | aRMS-5 | 688 | 175 | 385 | 298 | 319 |
|  | eRMS-1.1 | 420 | 115 | 203 | 79 | 176 |
|  | eRMS-1.2 | 1738 | 635 | 874 | 877 | 83 |
|  | eRMS-4 | 1348 | 1734 | 538 | 940 | 92 |
|  | eRMS-8.1 | 601 | 766 | 293 | 435 | 274 |
|  | eRMS-8.2 | 859 | 791 | 487 | 499 | 307 |
|  | eRMS-8.3 | 695 | 1293 | 355 | 475 | 121 |
|  | eRMS-2.1 | 572 | 126 | 192 | 117 | 321 |
|  | eRMS-2.2 | 1380 | 396 | 726 | 514 | 235 |
|  | eRMS-3.2 | 1502 | 895 | 856 | 497 | 616 |
| **Subtype** | aRMS | 9106 | 2193 | 4326 | 3507 | 2679 |
|  | eRMS | 9115 | 6751 | 4524 | 4433 | 2225 |
| **Treatment** | NA | 2246 | 429 | 1246 | 816 | 547 |
|  | off-treatment | 6394 | 3589 | 3058 | 3003 | 1284 |
|  | on-treatment | 9581 | 4926 | 4546 | 4121 | 3073 |
| **Cycling properties** | Cycling | 16675 | 928 | 8726 | 2588 | 1568 |
|  | Non-cycling | 1546 | 8016 | 124 | 5352 | 3336 |

**Extended Data Table 5:** PDC culture characteristics.

| PDX | Alternative name | Medium | bFGF/EGF | Passage # scRNAseq | Passage #CyTOF |
| --- | --- | --- | --- | --- | --- |
| aRMS-1 | IC-pPDX-104 | Complete F12 | + | 2 | 5 |
| aRMS-2 | IC-pPDX-29 | Complete NB | + | 6 | 10 |
| aRMS-3 | IC-pPDX-35 | Complete NB | + | 1 |  |
| aRMS-4 | SJRHB013759_X1 (Mast118) | Complete NB | + | 7 |  |
| aRMS-5 | Berlin 13304 | Complete NB | + | 14 |  |
| aRMS-6 | RMS-ZH-003 | Complete NB | + |  | 9 |
| aRMS-7 | Berlin 11410 |  |  |  |  |
| aRMS-8 | Berlin 14419 |  |  |  | 4 |
| aRMS-9 | Berlin 10752 |  |  |  | 5 |
| aRMS-10 | RMS-Wi-001B | Complete F12 | + |  |  |
| aRMS-11 | SJRHB010463_X16 (Mast60) |  |  |  |  |
| eRMS-1.1 | SJRHB011_Y (Rh70) | Complete NB | - | 5 | 8 |
| eRMS-1.2 | SJRHB011_X (Rh73) | Complete NB | - | 2 | 8 |
| eRMS-2.1 | SJRHB012_X (Rh71) | Complete NB | - | 4 | 3 |
| eRMS-2.2 | SJRHB012_Y (Rh74) | Complete NB | - | 3 |  |
| eRMS-2.3 | SJRHB012_Z (Rh75) | Complete NB | - |  |  |
| eRMS-3.1 | SJRHB13758_X1  (Mast111) |  |  |  |  |
| eRMS-3.2 | SJRHB13758_X2  (Mast139) | Complete NB | + | 4 |  |
| eRMS-4 | IC-pPDX-82 | Complete NB | - | 1 | 0 |
| eRMS-5 | Berlin 11410 |  |  |  |  |
| eRMS-6 | Berlin 12181 |  |  |  |  |
| eRMS-8.1 | Berlin 13454 | Complete NB | - | 5 |  |
| eRMS-8.2 | Berlin 13870 | Complete NB | - | 2 | 1 |
| eRMS-8.3 | Berlin 13933 | Complete NB | - | 4 | 7 |

| Sample | Hashtag | Barcode sequence | Sequencing run |
| --- | --- | --- | --- |
| KFR | Totalseq^TM^-B0251 hashtag 1 | GTCAACTCTTTAGCG | X |
| Rh4 | Totalseq^TM^-B0252 hashtag 2 | TGATGGCCTATTGGG |  |
| RMS | Totalseq^TM^-B0253 hashtag 3 | TTCCGCCTCTCTTTG |  |
| aRMS-1 | Totalseq^TM^-B0251 hashtag 1 | GTCAACTCTTTAGCG | X |
| aRMS-2 | Totalseq^TM^-B0252 hashtag 2 | TGATGGCCTATTGGG |  |
| aRMS-3 | Totalseq^TM^-B0253 hashtag 3 | TTCCGCCTCTCTTTG |  |
| aRMS-4 | Totalseq^TM^-B0251 hashtag 1 | GTCAACTCTTTAGCG | X |
| aRMS-5 | Totalseq^TM^-B0253 hashtag 3 | TTCCGCCTCTCTTTG |  |
| eRMS-1.1 | Totalseq^TM^-B0253 hashtag 3 | TTCCGCCTCTCTTTG | X |
| eRMS-1.2 | Totalseq^TM^-B0251 hashtag 1 | GTCAACTCTTTAGCG |  |
| eRMS-4 | Totalseq^TM^-B0253 hashtag 3 | TTCCGCCTCTCTTTG |  |
| eRMS-2.1 | Totalseq^TM^-B0253 hashtag 3 | TTCCGCCTCTCTTTG | X |
| eRMS-2.2 | Totalseq^TM^-B0251 hashtag 1 | GTCAACTCTTTAGCG |  |
| eRMS-3.2 | Totalseq^TM^-B0253 hashtag 3 | TTCCGCCTCTCTTTG |  |
| eRMS-8.1 | Totalseq^TM^-B0253 hashtag 3 | TTCCGCCTCTCTTTG | X |
| eRMS-8.2 | Totalseq^TM^-B0251 hashtag 1 | GTCAACTCTTTAGCG |  |
| eRMS-8.3 | Totalseq^TM^-B0253 hashtag 3 | TTCCGCCTCTCTTTG |  |
| Rh4 shSCR no DOX | TotalSeq™-B0251 Hashtag 1 | GTCAACTCTTTAGCG | X |
| Rh4 shSCR with DOX | TotalSeq™-B0252 Hashtag 2 | TGATGGCCTATTGGG |  |
| Rh4 shP3F1 no DOX | TotalSeq™-B0253 Hashtag 3 | TTCCGCCTCTCTTTG |  |
| Rh4 shP3F1 no DOX | TotalSeq™-B 0254 Hashtag 4 | AGTAAGTTCAGCGTA |  |
| KFR shSCR no DOX | TotalSeq™-B 0255 Hashtag 5 | AAGTATCGTTTCGCA |  |
| KFR shSCR with DOX | TotalSeq™-B 0258 Hashtag 8 | CTCCTCTGCAATTAC |  |
| KFR shP3F1 no DOX | TotalSeq™-B 0257 Hashtag 7 | TGTCTTTCCTGCCAG |  |
| KFR shP3F1 no DOX | TotalSeq™-B 0256 Hashtag 6 | GGTTGCCAGATGTCA |  |

**Extended Data Table 6:** Totalseq^TM^-B hashtag antibody list used for staining.

X, samples were pooled together in the same 10X run.
