## Extended Data Figures for "Single-cell mapping of tumor heterogeneity in pediatric rhabdomyosarcoma reveals developmental signatures with therapeutic relevance"

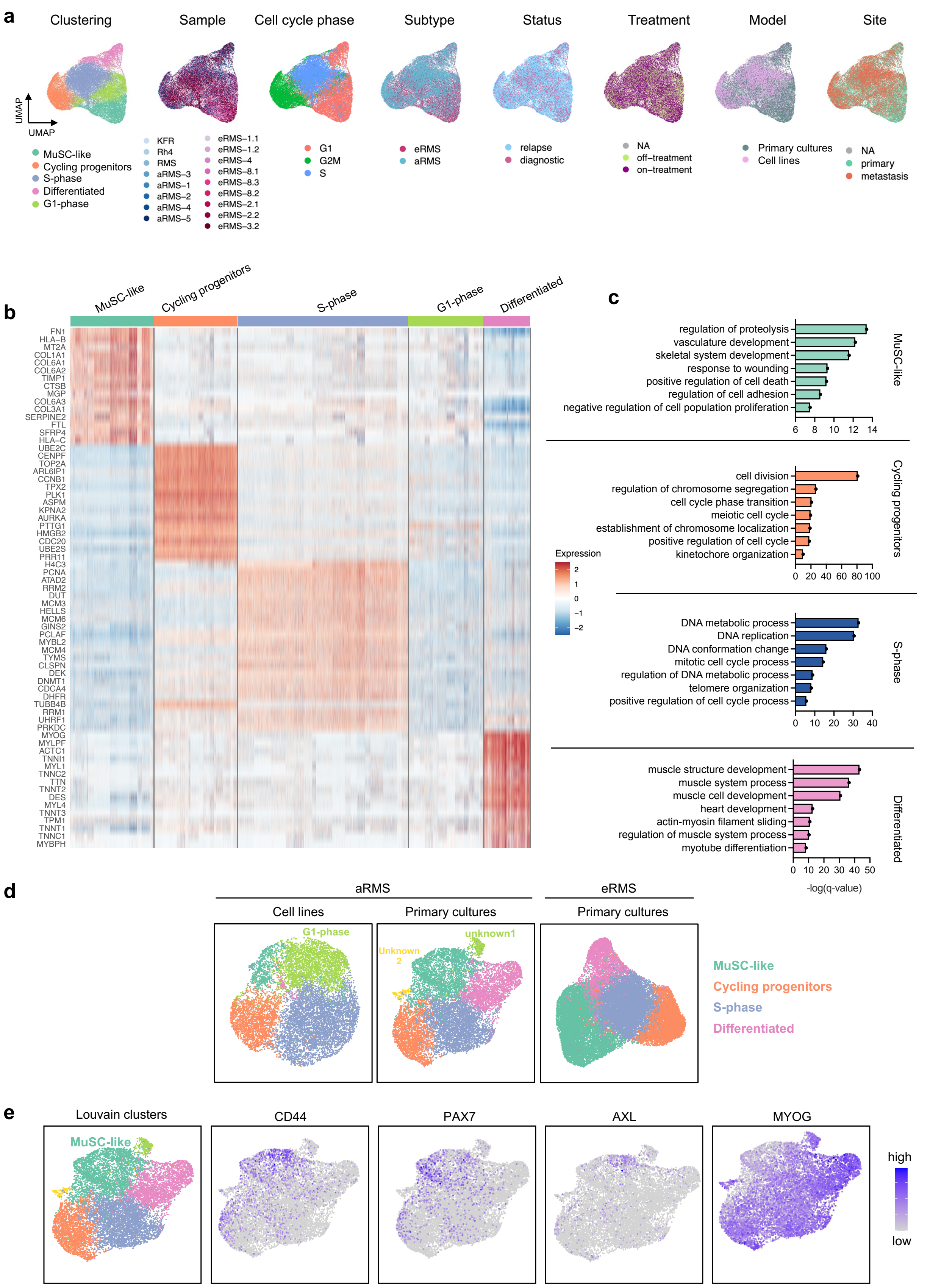


**Extended Data Figure 1: RMS single-cell gene signatures**

**a.** UMAP plot of 48,859 cells after integration. Cells are colored based on the populations identified by Louvain clustering, the sample of origin, the inferred cell cycle status, the RMS subtype, the status, treatment and site of the tumor at the time of PDX generation and on the model. **b.** Heatmap of the top 15 gene markers for each cluster (top bars) across the combined RMS dataset. Relative $\log_2$ fold change of gene expression (color bar) is shown **c.** Top seven biological processes enriched in each cluster calculated by Metascape (Zhou et al., *Nature Communications,* 2019). **d.** UMAP plots of combined aRMS primary cultures or cell lines and of eRMS primary cultures after removing inter-sample differences by SCT algorithm. The identified cellular states are indicated. **e.** UMAP plots of combined aRMS primary cultures colored based on the identified Louvain clusters (left panel) or based on expression of CD44, PAX7, AXL and MYOG.

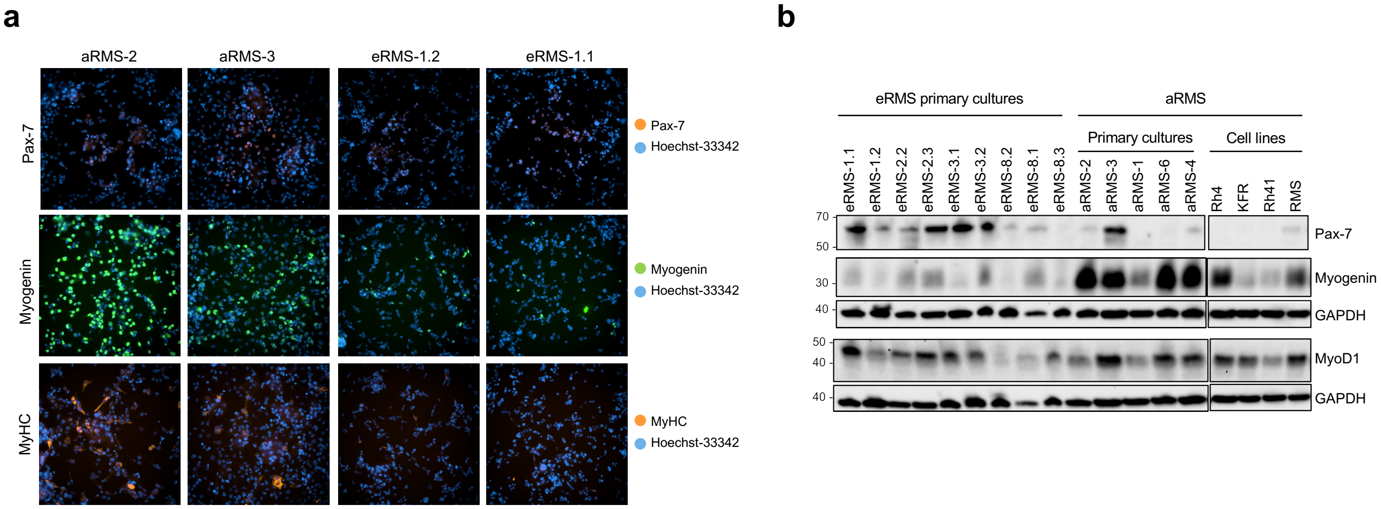


**Extended Data Figure 2: aRMS are skewed towards later differentiation stages than eRMS tumors**

**a.** Representative immunofluorescence images of aRMS and eRMS primary cultures stained for Pax-7, myogenin or MyHC.

**b.** Representative western blots of Pax-7, MyoD and myogenin protein expression in a panel of RMS primary cultures and cell lines. GAPDH was used as a loading control.

*, $P < 0.05$; **, $P < 0.01$.

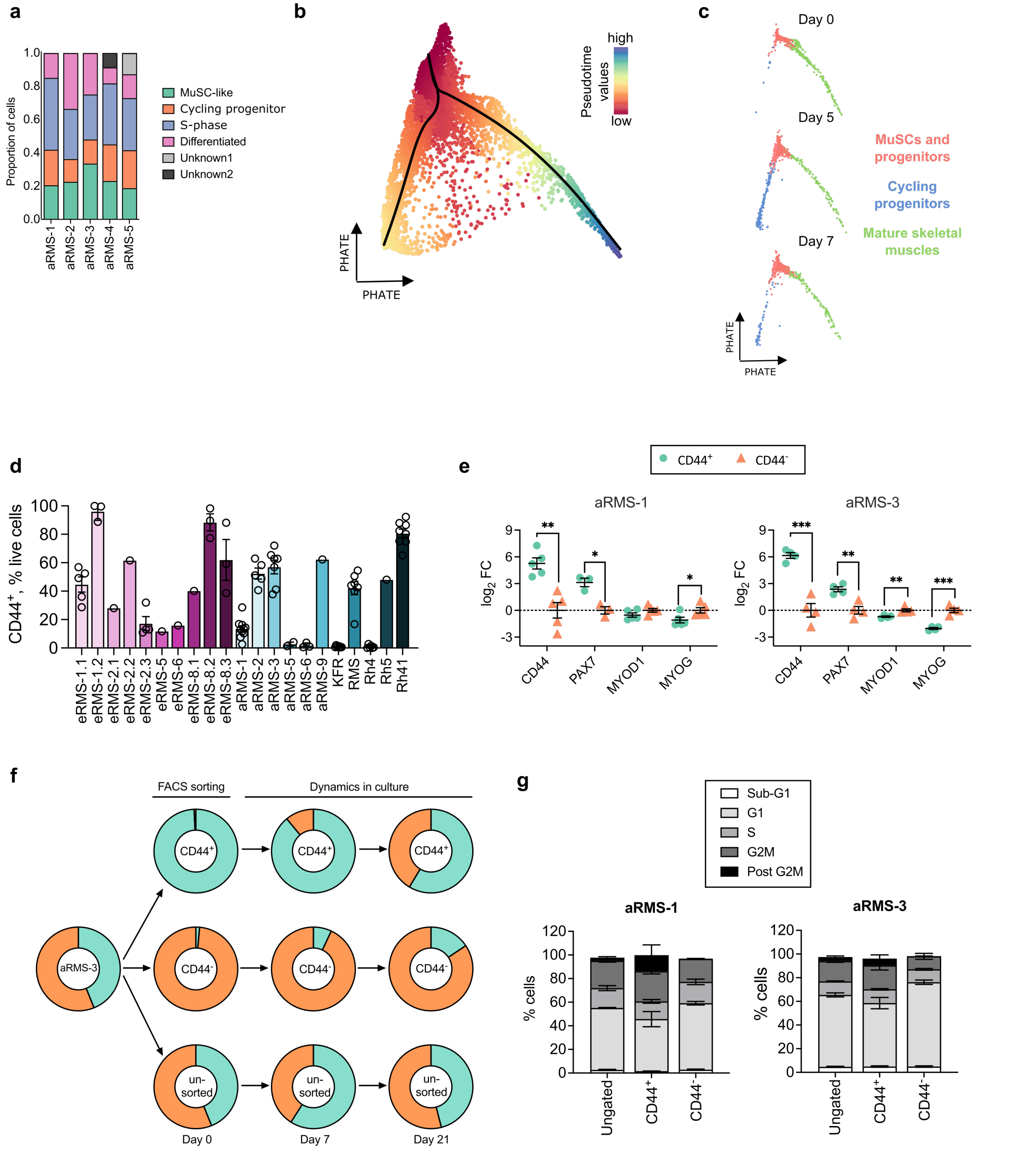


**Extended Data Figure 3. Hierarchical organization of RMS tumors**

**a.** Fraction of cells belonging to the identified clusters across $n$ = 5 aRMS primary cultures after integration using SCT-correction. Only clusters that were commonly shared across different cultures ("MuSC-like", "Cycling progenitor", "S-phase", "Differentiated") were selected for pseudotime analysis. The clusters "Unknown1" and "Unknown2" were only found in the sample aRMS-5, respectively aRMS-4. For this reason, both clusters were excluded for PHATE dimensionality reduction and pseudotime trajectory calculation. **b.** PHATE plots showing the pseudotime trajectories of the combined aRMS primary cultures/mouse scRNAseq (De Micheli et al., *Cell Reports*, 2020) datasets. Trajectories were calculated using the *Slingshot* package (*MuSC-like* cells were set as the trajectory start); black lines represent the inferred trajectories whereas cells are colored based on calculated pseudotime values. **c.** PHATE plots of mouse muscle scRNAseq dataset (De Micheli et al., *Cell Reports*, 2020) at different days (0, 5, 7) after muscle injury. Cells are colored based on the subpopulations identified in the original publication. **d.** CD44 cell surface expression measured by flow cytometry in a panel of primary cultures and cell lines of eRMS (pink) or aRMS (blue) subtypes. Data are represented as mean ± SEM of the indicated number of biological replicates. **e.** qRT-PCR data generated from FACS-sorted $CD44^+$ and $CD44^-$ subpopulations. $Log_2$ fold change mRNA levels of $CD44^+$ normalized to $CD44^-$ are depicted. Data are represented as mean ± SEM of $n \geq 4$ biological replicates; multiple unpaired *t*-tests. **f.** Flow cytometry analysis of the stability of $CD44^+$ and $CD44^-$ subpopulations in aRMS-3 cells. Unsorted reference is also shown. Data are represented as mean ± SEM of $n \geq 2$ biological replicates. **g.** Cell cycle distribution of $CD44^+$, $CD44^-$ and ungated subpopulations. Data are represented as mean ± SEM of $n \geq 2$ biological replicates.
*, $P < 0.05$; **, $P < 0.01$; ***, $P < 0.001$

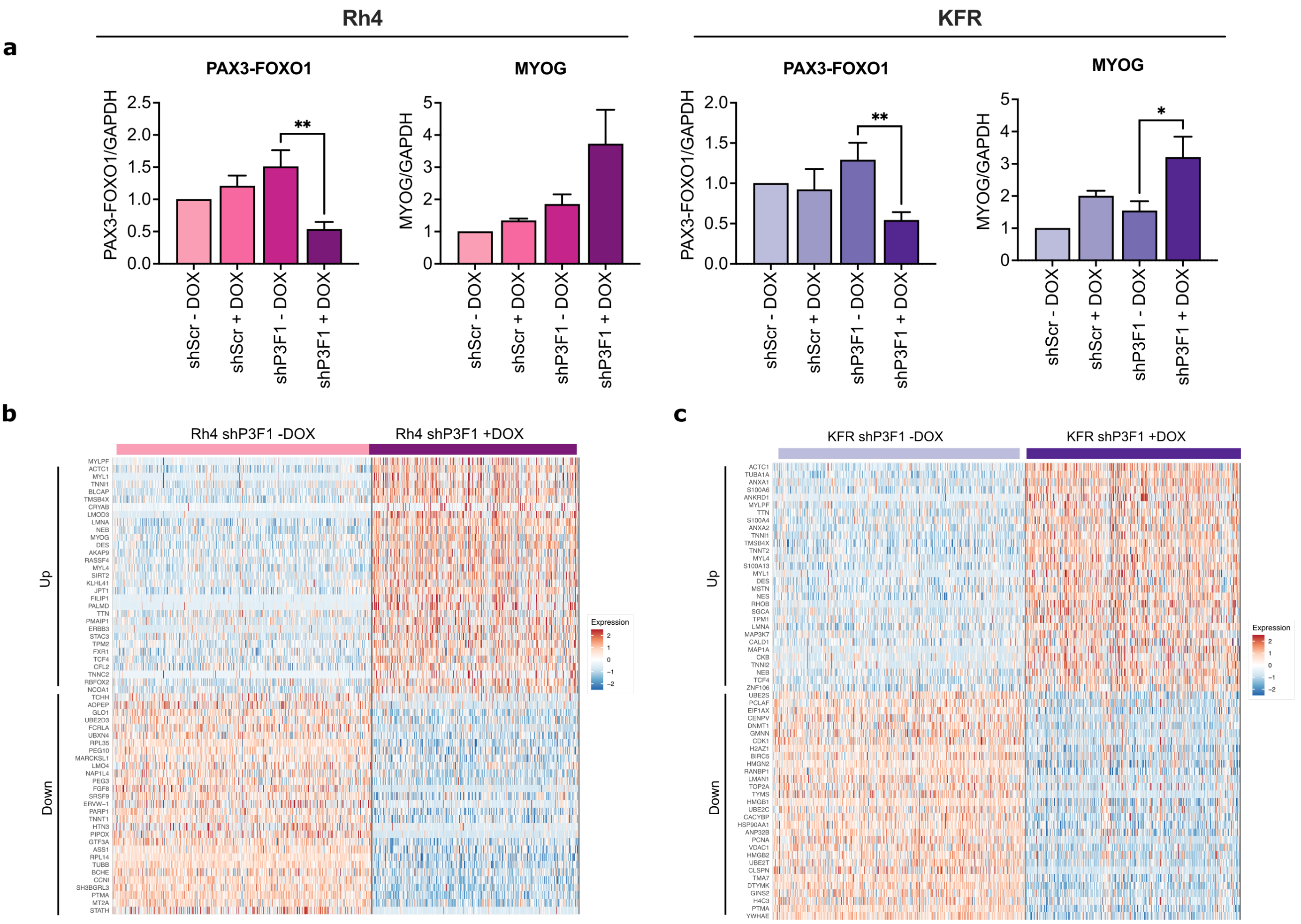


**Extended Data Figure 4. Single-cell responses upon P3F1 downregulation**

**a.** Western blot quantification of Rh4 or KFR cells treated with or without DOX for 48 hrs. Data are represented as mean ± SEM of $n \geq 5$ biological replicates; ordinary one-way ANOVA with Dunnet's multiple comparison test. **b and c.** Heatmap plots showing log fold change of the top 30 genes differentially expressed in Rh4 (**b**) or KFR (**c**) cells transduced with shP3F1 construct upon DOX treatment. Relative $\log_2$ fold change of gene expression (color bar) is shown.

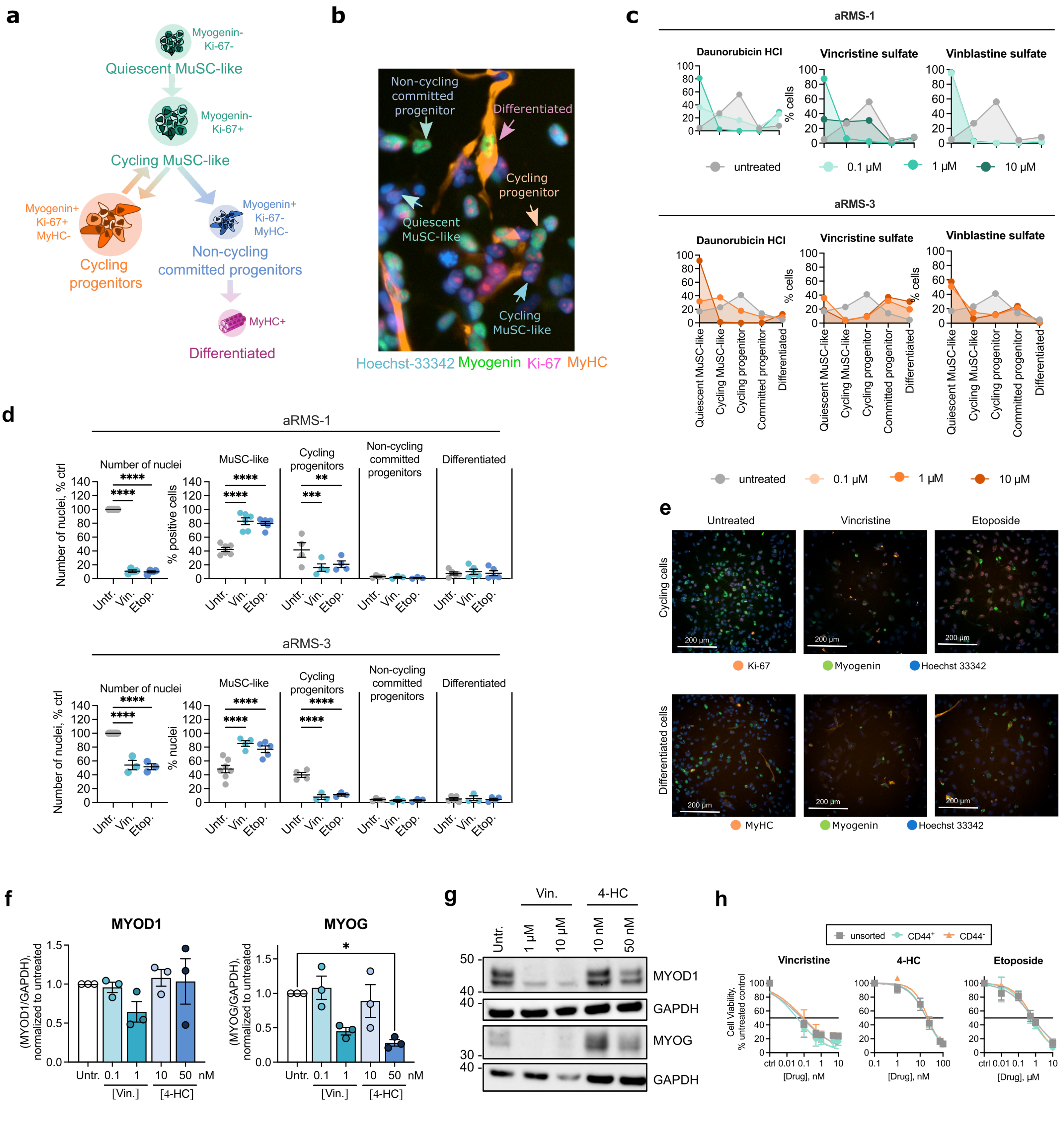


**Extended Data Figure 5: Chemotherapy shifts aRMS cells towards progenitor states**

**a.** Myogenin, Ki-67 and MyHC marker positivity across the aRMS cellular states identified by single-cell analysis. **b.** Representative immunofluorescence analysis of aRMS-1 cells. Cells representative for the indicated cellular states are marked with arrows. **c.** Cellular shifts upon treatment with the indicated drugs in aRMS-1 and aRMS-3 cells. Grey lines represent basal cellular composition in untreated controls. **d.** IF quantification of aRMS-1 or aRMS-3 cells exposed to vincristine sulfate (10 nM in aRMS-1, 10 µM in aRMS-3 cells) or etoposide (1 µM in aRMS-1, 10 µM in aRMS-3 cells) for 72 hrs. Cell subpopulations were defined as follows: *MuSC-like* (myogenin$^-$); *cycling progenitors* (myogenin$^+$ Ki-67$^+$); *non-cycling committed progenitors* (myogenin$^+$ Ki-67$^-$), *differentiated* (MyHC$^+$). Data are represented as mean ± SEM of the indicated number of biological replicates; ordinary two-way ANOVA with uncorrected Fisher's LSD. **e.** Representative images of aRMS-3 cells exposed to 1 µM vincristine sulfate or to 10 µM etoposide for 72 hrs. **f.** WB quantification of aRMS-1 cells exposed to vincristine or to 4-HC for 48 hrs at the indicated concentrations. Data are represented as mean ± SEM of $n$ = 3 biological replicates; ordinary two-way ANOVA with Dunnett's multiple comparison test. **g.** WB analysis of aRMS-3 cells exposed to vincristine or to 4-HC for 48 hrs at the indicated concentrations. **h.** Dose-response curves of vincristine sulfate, 4-HC and etoposide in FACS-sorted aRMS-1 subpopulations.
*Untr.*: untreated; *Vin.*: vincristine sulfate; *Etop.*: etoposide; *4-HC*: 4-hydroperoxycyclophosphamide.
*, $P < 0.05$; **, $P < 0.01$; ***, $P < 0.001$.; ****, $P < 0.0001$.

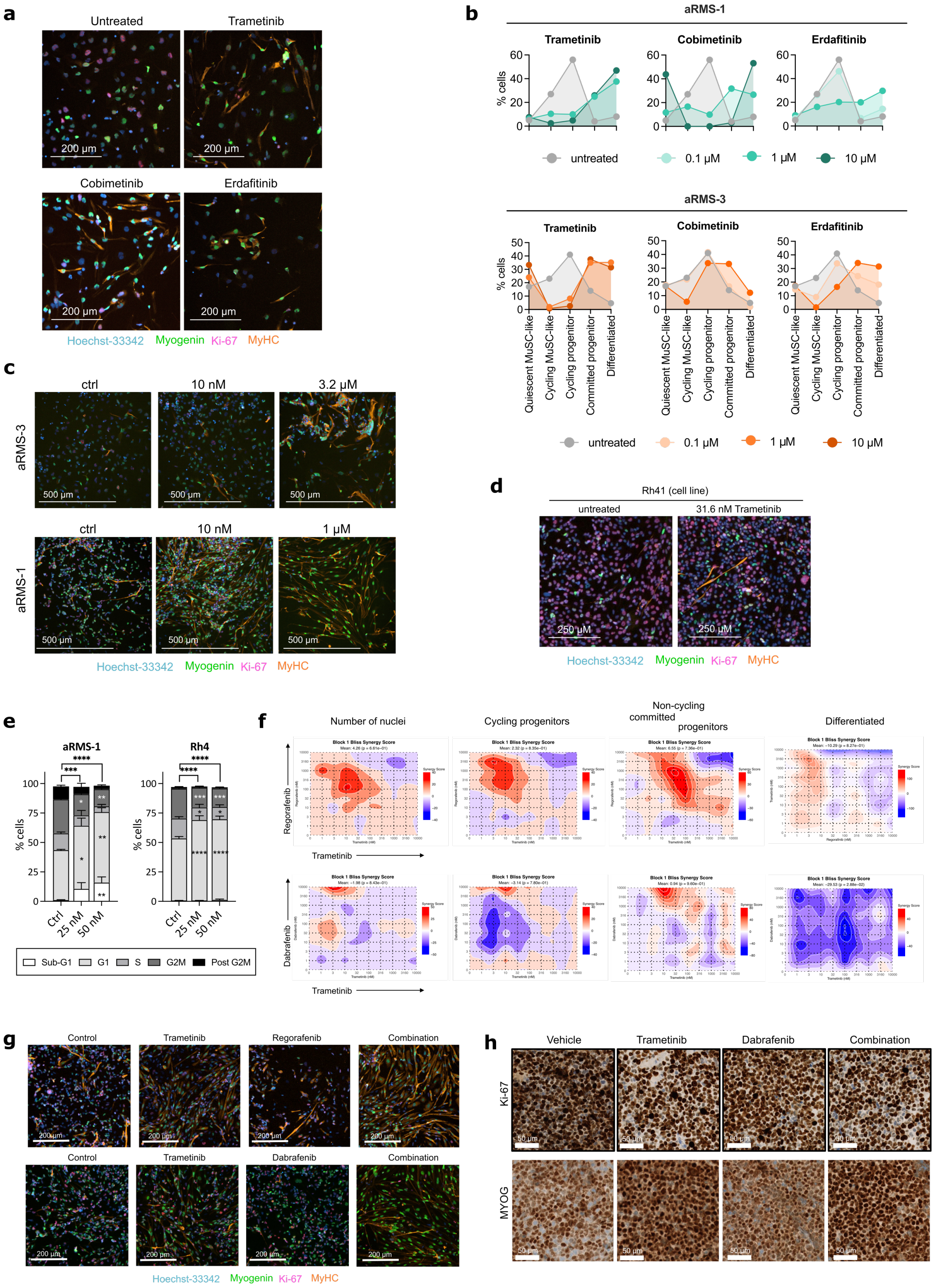


**Extended Data Figure 6: Trametinib effect on aRMS primary cultures and cell lines**

**a.** Representative immunofluorescence images of aRMS-3 cells following 72 hrs drug treatment at 1 μM. **b.** Cellular shifts upon treatment with differentiating drug hits trametinib, cobimetinib or erdafitinib in aRMS-1 and aRMS-3 cells at the indicated concentrations. Grey lines represent basal cellular composition in untreated controls. **c and d.** Representative IF images of aRMS primary cultures (**c**) or of the cell line Rh41 (**d**) exposed to the indicated concentrations of trametinib for 72 hrs. **e.** Quantification of cell cycle phases after exposure of aRMS-1 or Rh4 cells to the indicated concentrations of Trametinib for 96 hrs. Data are represented as mean ± SEM of $n \geq 3$ biological replicates; ordinary two-way ANOVA with Dunnet's multiple comparison test. **f.** Synergy maps of trametinib-regorafenib (top row) or trametinib-dabrafenib (bottom row) combinations. Cells from aRMS-1 were exposed to the drugs for 72 hrs and then processed for MYOscopy. Synergy was calculated according to the Bliss model (Bliss, CI. *Annals of Applied Biology*, 1939) on SynergyFinder (Zheng, S et al. *Genomics Proteomics Bioinformatics*, 2022). **g.** Representative immunofluorescence images of aRMS-1 cells exposed to 10 nM trametinib, 10 μM dabrafenib, 1 μM regorafenib or to the combination of trametinib-regorafenib (top row) or trametinib-dabrafenib (bottom row), measured by MYOscopy. Data are represented as mean ± SEM of $n = 4$ biological replicates; ordinary one-way ANOVA with Tukey's multiple comparison test. **h.** Expression of Ki-67 and myogenin (MYOG) as determined by immunohistochemistry in aRMS-1 PDX tumors following *in vivo* treatment with 1 mg/kg trametinib, 15 mg/kg dabrafenib or with their combination.

*, $P < 0.05$; **, $P < 0.01$; ***, $P < 0.001$.; ****, $P < 0.0001$.
